# Amyloid β Induces Lipid Droplet-Mediated Microglial Dysfunction in Alzheimer’s Disease

**DOI:** 10.1101/2023.06.04.543525

**Authors:** Priya Prakash, Palak Manchanda, Evi Paouri, Kanchan Bisht, Kaushik Sharma, Jitika Rajpoot, Victoria Wendt, Ahad Hossain, Prageeth R. Wijewardhane, Caitlin E. Randolph, Yihao Chen, Sarah Stanko, Nadia Gasmi, Anxhela Gjojdeshi, Sophie Card, Jonathan Fine, Krupal P. Jethava, Matthew G. Clark, Bin Dong, Seohee Ma, Alexis Crockett, Elizabeth A. Thayer, Marlo Nicolas, Ryann Davis, Dhruv Hardikar, Daniela Allende, Richard A. Prayson, Chi Zhang, Dimitrios Davalos, Gaurav Chopra

## Abstract

Several microglia-expressed genes have emerged as top risk variants for Alzheimer’s disease (AD). Impaired microglial phagocytosis is one of the main proposed outcomes by which these AD-risk genes may contribute to neurodegeneration, but the mechanisms translating genetic association to cellular dysfunction remain unknown. Here we show that microglia form lipid droplets (LDs) upon exposure to amyloid-beta (Aβ), and that their LD load increases with proximity to amyloid plaques in brains from human patients and the AD mouse model 5xFAD. LD formation is dependent on age and disease progression and is prominent in the hippocampus in mice and humans. Despite differences in microglial LD load between brain regions and sexes in mice, LD-laden microglia exhibited a deficit in Aβ phagocytosis. Unbiased lipidomic analysis identified a decrease in free fatty acids (FFAs) and a parallel increase in triacylglycerols (TGs) as the key metabolic transition underlying LD formation. DGAT2, a key enzyme for converting FFAs to TGs, promotes microglial LD formation and is increased in 5xFAD and human AD brains. Inhibition or degradation of DGAT2 improved microglial uptake of Aβ and drastically reduced plaque load in 5xFAD mice, respectively. These findings identify a new lipid-mediated mechanism underlying microglial dysfunction that could become a novel therapeutic target for AD.

## Introduction

Alzheimer’s disease (AD) is a progressive neurodegenerative disorder of the aging human population. Accumulation of amyloid-beta (Αβ) is a defining histological hallmark of the AD brain^1^. However, failures in Αβ-targeted clinical trials^2^, combined with findings showing no apparent correlation between cognitive decline and overall plaque load in AD patients^3^, suggest that additional mechanisms may be crucially involved in AD etiopathogenesis. Genome-wide association studies (GWAS) identified many AD risk genes related to the immune response and microglia, including the phagocytic receptors CD33 and TREM2^4,5,6,7,8^. Single-cell RNA sequencing identified gene signatures characteristic of prominent protective and dysfunctional microglial subpopulations in AD^9,10^. Moreover, a study that focused on human brain disease-associated variants of non-coding regulatory regions in a cell-specific manner identified multiple sporadic AD risk variants specifically in microglial transcriptional enhancers^11^, further highlighting the involvement of microglia in AD pathogenesis.

In the AD brain, reactive microglia cluster around Aβ plaques^12^ and form a physical barrier believed to restrict plaque propagation^13,14^. During early stages of AD, microglial reactivity is considered as beneficial for Αβ clearance^15^; however, sustained inflammation likely contributes to neurodegeneration^12^. Increased pro-inflammatory gene expression in response to accumulating Αβ in older AD mice leads to decreased microglial Αβ clearance receptors or Αβ-degrading enzymes, thereby promoting further Αβ accumulation^16^. Furthermore, the ability of microglia to remove Αβ declines over time, supporting that the progression of amyloid pathology correlates with impaired capacity of microglia to phagocytose Αβ^17,16^.

In addition to genes and proteins, changes in microglial lipid content can also affect their state and function^18,19,20,21^. Cellular lipids regulate functions like migration and phagocytosis^22^, are important for immune cell modulation and signaling^23^, and their dysregulation has been linked to neurodegenerative disorders, including AD^24^. Top AD-risk genes such as TREM2, APOE, and INPP5D are directly related to lipid metabolism^25,26,27^. Although lipid droplets (LDs, first described as fat particles by Alois Alzheimer^28^) were initially considered to be passive fat deposits^29^, they are dynamic intracellular organelles (diameter <1–100 μm) that regulate lipid metabolism. LDs consist of a phospholipid monolayer containing a core of neutral lipids like triacylglycerols (TAGs) and cholesteryl esters (CEs)^30^. Inflammatory triggers like lipopolysaccharide (LPS)^31,19^, fatty acids^32^, and aging^20^ can result in LD accumulation in primary microglial cells and cell lines. Further, human iPSC-derived microglia were enriched in LDs in a chimeric mouse model of AD^33^. However, it is not known if Aβ can directly induce LD formation in microglia, and if/how changes to their lipid or metabolite composition can affect microglial function in AD.

Here we show that plaque-associated microglia closely associated with Aβ plaques contained more LDs and had larger cell bodies and shorter processes in both mouse and human brains, highlighting a unique LD-laden microglial subtype in AD. Extensive lipidomic and metabolomic profiling in microglia revealed specific types of lipids and metabolic pathways likely responsible for their LD-laden phenotype in the 5xFAD mouse model, which was more pronounced in microglia from females compared to males. Functionally, LD-laden microglia showed reduced phagocytosis of Aβ compared to age-matched WT microglia. Mechanistically, we found that Aβ treatment alone promotes a drastic shift in lipid content in microglia isolated from WT brains, even within only 24 hours of Aβ exposure. Extensive lipidomic characterization of these microglia revealed a gradual increase in TAG content following Aβ treatment, which was similar to the lipid composition changes we detected in microglia from 5xFAD mice. Based upon this finding, we identified diacylglycerol O-acyltransferase 2 (DGAT2), a key enzyme for the conversion of FFAs to TAGs, as also being the key catalyst for Aβ-induced LD formation in microglia. DGAT2 protein expression was increased in both mouse and human AD brains, and inhibiting DGAT2 increased the phagocytosis of Aβ by 5xFAD microglia. Importantly, degradation of the DGAT2 enzyme in the brain of 5xFAD mice drastically reduced amyloid plaque load. Our study thus highlights DGAT2 as a promising target for preventing or reversing phagocytic dysfunction in LD-laden microglia in the AD brain.

### Microglia accumulate lipid droplets in an age-, sex-, and region-dependent manner in the 5xFAD model of Alzheimer’s disease

Prior literature has linked the accumulation of LDs in microglia with inflammation, aging, and impaired phagocytosis, all of which are also hallmarks of AD^31,20,33^. We asked if/how exposure to Aβ could induce LD formation in microglia, and/or alter their lipid or metabolite composition in a way that affects their function in AD. We used a mouse model with five familial AD mutations (5xFAD)^34^, which progressively accumulates extensive Aβ plaques (especially in the subiculum and deep cortical layers) starting around 2-3 months of age. 5xFAD mice also develop prominent gliosis, inflammation, neuronal loss, behavioral impairments, and sex-distinct systemic metabolic changes^35,34,36^. We isolated microglia from 5xFAD and WT mice and performed flow cytometry analysis after staining for their neutral lipids (**Fig. 1a; Fig. S1a**). Although there was no difference in younger mice (**Fig. 1b**), microglia from 5-7-month-old female 5xFAD mice showed significantly higher LD content (1.58-fold) compared to cells from age-matched controls. Among LD^+^ cells, almost all 5xFAD microglia had high LD content, whereas WT microglia had lower LD content (**Fig. 1c**). Microglia from 5-7-month-old male 5xFAD mice also showed a significant but modest increase (1.15-fold) in LDs compared to WT (**Fig. 1d**), however a higher number of LD^+^ microglia had high LD content in cells from 5xFAD compared to WT male mice (**Fig. 1e**). These findings demonstrate that microglia show increased LD load, in a mouse model of AD with extensive Aβ brain deposition, albeit with some age- and sex-dependent variability.

**Fig. 1:**
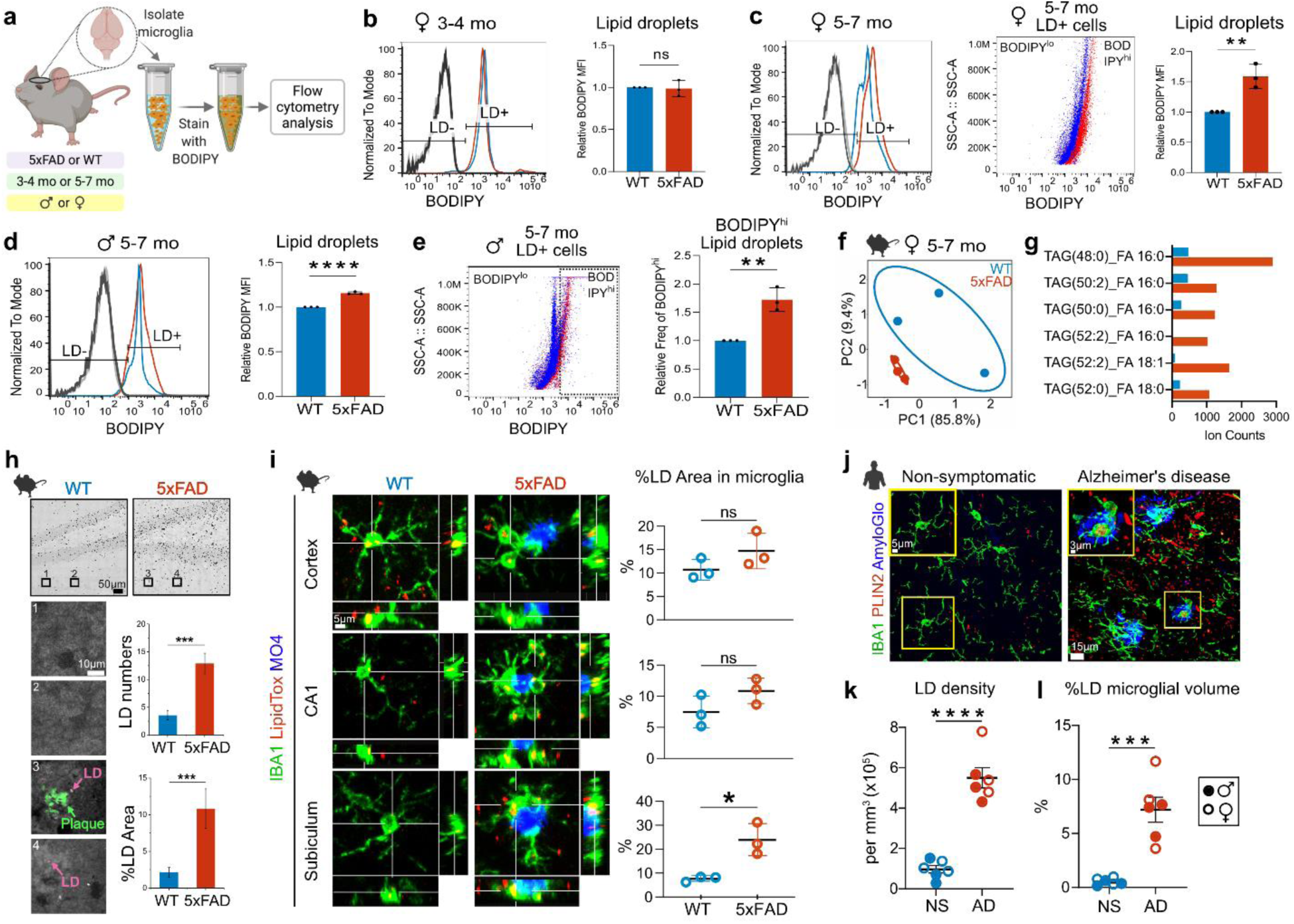
LD abundance in microglia is age-, sex-, and region-dependent in the AD brain. **a.** Experimental design for labeling and quantifying LDs in acutely-isolated microglia from 5xFAD and WT male and female mice at 3-4 or 5-7 months old. LDs were labeled with BODIPY dye and quantified using flow cytometry. **b.** Representative graph (left) and quantification (right) of median fluorescence intensity (MFI) of LDs in live microglia (CD11b^+^DAPI^-^) from 3-4-month-old female mice show no increase in LD content in cells from 5xFAD (red line and bar) compared to WT animals (blue line and bar). Data represent mean ± SD. Unpaired t-test, N=3 separate experiments, each including one WT and one 5xFAD mouse. **c.** Quantification of LDs in microglia from 5-7-month-old female mice shows an increase (shift towards higher BODIPY fluorescence intensity) in LDs from 5xFAD (red) compared to WT (blue) microglia. Dot plot shows a homogeneous population of BODIPY^hi^ microglia within the LD^+^ cell subset from 5xFAD mice (red dots) compared to microglia from WT mice (blue dots); \*\**P*= 0.0068. **d.** Quantification of LDs in microglia from 5-7-month-old male mice shows an increase in LD content in 5xFAD microglia compared to WT; \*\*\*\**P*= 0.00003. **e.** Comparison between BODIPY^hi^ and BODIPY^lo^ cell populations within LD^+^ microglia from 5-7-month-old male mice shows more BODIPY^hi^ microglial cells in 5xFAD mice (red dots) compared to microglia from WT mice (blue dots). Quantification shows the relative frequency of microglia containing LDs in the BODIPY^hi^ gate in 5xFAD and WT microglia; *\*\*P*= 0.0039. For **c-e**: Data represent mean ± SD. Unpaired t-test, N=3 separate experiments, each including two WT and two 5xFAD mice. **f.** Principal component analysis (PCA) plot depicts a clear separation based upon variation in microglial lipidomes from 5-7-month-old WT and 5xFAD female mice. **g.** Graph shows the identification and relative amounts of specific TAG lipid species that were increased in microglia from 5-7-month-old female 5xFAD mice compared to cells from WT controls. For **f-g:** N=3 separate experiments, each from one WT and one 5xFAD mouse. **h.** Label-free SRS imaging of LDs and Aβ plaques in 5xFAD and WT brain hippocampal slices. Increased LDs were observed and quantified in 5xFAD brain sections, often associated with Aβ plaques (3, 4), compared to WT tissues (1, 2). Twelve areas were quantified for each group (WT and 5xFAD), from the same brain section. Unpaired t-test, N=2 separate experiments, each including one WT and one 5xFAD mouse. **i.** Immunofluorescence of IBA1, and counter-staining for LDs (LipidTox), and Aβ plaques (Methoxy XO4; MO4) in the cortex, CA1, and subiculum regions from 5xFAD and WT mice. Quantification showed a trend towards increased % LD area within microglia in the cortex and CA1 regions, which was statistically significant in the subiculum (\**P=* 0.0147) from 5xFAD compared to WT mice. Data represent mean ± SD. Unpaired t-test, N=3 separate experiments, each including one WT and one 5xFAD mouse. **j.** Detection of lipid droplets in human hippocampal formalin-fixed paraffin-embedded (FFPE) tissue from AD and non-symptomatic (NS) cases. Immunofluorescence was performed on 15µm sections for the detection of lipid droplets (PLIN2), Aβ plaques (AmyloGlo), and microglia (IBA1). Representative images show an increase in the density of PLIN2^+^ LDs in AD compared to non-symptomatic cases. Higher magnification inserts show an increase of LDs in microglia from AD patients compared to NS individuals. **k.** Quantification of the number of PLIN2^+^ LDs per mm^3^ of imaged volume (LD density) shows a significant increase in AD compared to NS cases; \*\*\*\**P*= 0.000007. **l.** Quantification of percentage of microglial volume occupied by LDs over the total microglial volume per imaged volume of hippocampal tissue shows an increase in AD compared to NS cases; \*\*\**P*= 0.0002. For **k**, **l**: quantification was performed in 3D reconstructed confocal z-stacks using Imaris; Data represent mean ± SEM. Unpaired t-test, N=6 (3 males and 3 females) per group.

Next, we investigated if acutely-isolated microglia from 5xFAD mice showed alterations in their global lipid and metabolite profiles as they experienced a drastic increase in Aβ plaque load with age. We performed unbiased mass spectrometry lipidomic and metabolomic profiling using a modified multiple reaction monitoring (MRM)^37^ approach that allowed us to screen 1380 lipid species, categorized into 10 main classes (**Fig. S1b**), and over 700 metabolites. Such broad coverage and depth of profiling allowed us to explore possible differences in the lipid profiles of microglia from 5-7-month-old male and female 5xFAD mice and compare them to those from age- and sex-matched WT microglia in an unbiased manner (**Supplementary Tables ST1-ST2**). Female 5xFAD microglia showed clearly distinct profiles, (indicated by their clear separation in the principal component analysis (PCA) space, accounting for over 95% of the variation in the data) (**Fig. 1f**), with 105 differentially-regulated lipids that were primarily downregulated, compared to cells from WT controls **(Fig. S1c-e)**.

We further evaluated triacylglycerol (TAG) profiles in female microglia and found an increase in several long-chain species, including TAG(48:0), TAG(50:2), TAG(50:0), TAG(52:2), TAG(52:0) in 5xFAD compared to WT cells (**Fig. 1g, Supplementary Table ST3**). In contrast, the male lipidome showed less variability (poor separation in the PCA space) (**Fig. S1f**), in agreement with the more limited variation in LD content in male microglia when analyzed at the single-cell level (**Fig. 1d**). We also compared microglial metabolite profiles from the same tissues and found them to be distinctly different (**Fig. S2a**). Metabolites like fructose 6-phosphate and lactose were downregulated in 5xFAD male and female microglia compared to cells from WT controls, affecting several key metabolic pathways such as the citrate TCA cycle, glutamine and glutamate metabolism, arginine biosynthesis, etc. (**Fig S2b-c, Supplementary Tables ST4, ST5**). These results illustrate that following prolonged Aβ exposure, microglia exhibit: i) a unique lipid signature that likely facilitates the formation and accumulation of LDs within them in a sex-distinct manner, and ii) dysregulation in several key metabolic pathways, that could also impact their functions in the context of AD.

### Microglia have increased LD content in the hippocampus of 5xFAD mice and AD patients

We next asked if/how the increased LD content and related metabolic changes related to brain amyloid deposition, one of the prominent pathological hallmarks of 5xFAD mice and AD brains. We examined the spatial distribution of LDs by performing label-free stimulated Raman scattering (SRS) microscopy^38^ in hippocampal brain slices of 5xFAD mice. We found a significantly higher number of LDs, and a higher percentage of LD area overall, in chronic 5xFAD compared to age-matched WT hippocampi (**Fig. 1h**). Interestingly, some LDs appeared in close proximity to Aβ plaques in the 5xFAD tissue, as identified by their respective spectral signatures (**Fig. S3**).

To further profile the spatial distribution of LD-laden microglia relative to amyloid plaques, we combined histological staining with methoxy O4 (amyloid plaques) and LipidTox (lipids) with immunohistochemistry for IBA1 (microglia) in 5-7-month-old female 5xFAD brain sections. Even though aged WT brains had a high number of LDs in microglia (as also previously reported^20^), and in other cells (**Fig. 1i; S4a**), 5xFAD brains had significantly higher overall LD density and total LD area in cortex and parts of the hippocampus (CA1 and subiculum) compared to WT (**Fig. S4b-c**). The mean cell body area of LD**^+^** microglia was not significantly different than that of LD**^−^** cells (**Fig. S4d**), nor was the mean cell body area of LD**^+^** microglia in 5-7-month-old female 5xFAD mice compared to LD**^+^** cells in age and sex-matched WT controls (**Fig. S4e**). Similarly, the overall proportion of LD**^+^** microglia was similar across cortex, CA1, and subiculum (**Fig. S4f**), but the proportion of LD area within microglia was significantly increased in the subiculum of 5xFAD mice (**Fig. 1j**), indicating a preferential increase in LD load within microglia in this brain region.

Since the hippocampus is significantly affected by amyloid pathology in AD patients^39^, we also stained postmortem human hippocampal sections from non-symptomatic (NS) and AD patients for Aβ plaques, LDs, and microglia (**Fig. 1k**). After 3D reconstruction and segmentation of each stained structure imaged by high-resolution confocal microscopy, we performed volumetric co-localization analysis using Imaris (**Fig. S5)**. We measured significantly higher (5.7-fold increase) overall LD density (**Fig. 1l; Fig. S6)**, and—similar to the mouse model findings—a significantly higher percentage of microglial volume occupied by LDs in the hippocampus of AD patients compared to NS controls in both males and females (**Fig. 1m**).

### Chronic exposure of microglia to Aβ plaques promotes LD accumulation, alters microglial morphology, and impairs their phagocytic ability

We next examined the spatial relationship between plaque load and LD density and found them to be positively correlated in the cortex, CA1, and subiculum (**Fig. 2a**). Interestingly, the vast majority of plaque-proximal microglia were laden with LDs in all three brain regions (**Fig. 2b; Fig. S7a**). Particularly in the subiculum, plaque-proximal microglia demonstrated an amoeboid morphology, while plaque-distant cells typically had smaller cell bodies and longer processes (**Fig. 2c-d; Fig. S7b, c**). Importantly, this also translated in human microglia in the hippocampus of AD patients (both male and female), where plaque-proximal microglia had a significantly higher number of LDs than plaque-distant microglia, which decreased with distance from the nearest plaque (**Fig. 2e, f; Fig. S7d; Supplementary Movie 1**). In addition, LD**^+^** microglia closer to plaques (0-10μm) contained larger LDs, and their total LD load also decreased progressively with distance from the nearest plaque (**Fig. 2g; Fig. S7e**). The methodological approach implemented in Imaris to identify PLIN2**^+^**IBA1**^+^** cells and their LD load as a function of distance from the nearest plaque is schematically represented in **Fig S8**. Overall, these results suggest that a morphologically-distinct plaque-associated microglial phenotype characterized by larger LD load exists in both the human AD brain and the amyloid-rich 5xFAD animal model.

**Fig. 2:**
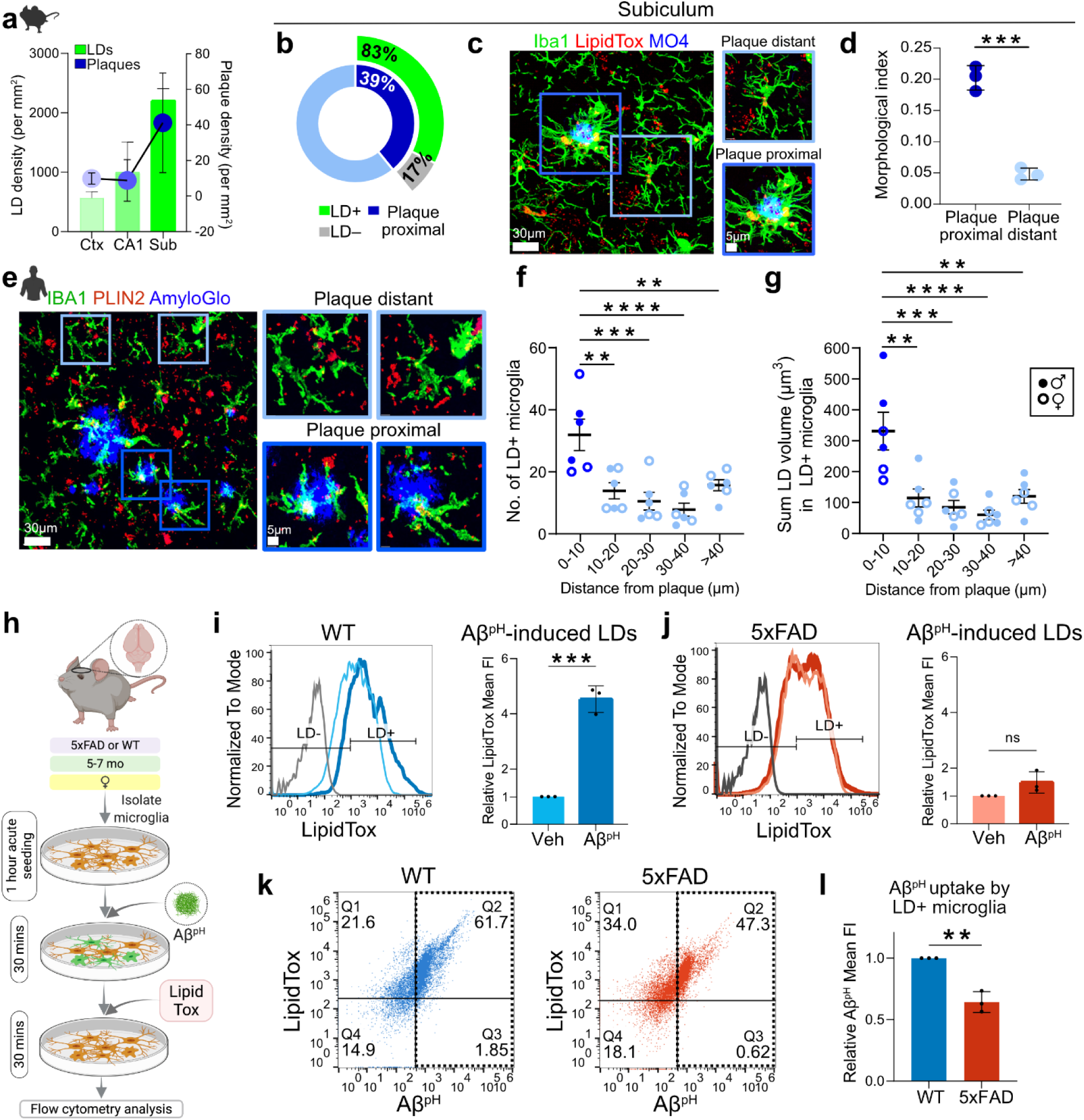
LD-laden microglia are in close proximity to amyloid plaques in mice and humans and exhibit phagocytosis deficits. **a.** Parallel quantification in three different 5xFAD brain regions, shows that LD density seems to correlate with plaque density, with the subiculum area of the hippocampus (Sub) where plaque density is highest also demonstrating the highest LD density compared to CA1 or cortex (Ctx). Data represent mean ± SD. N=3 5xFAD mice. **b.** Quantification of % LD^+^ microglia that are plaque-proximal or -distant in the subiculum of 5xFAD mice. Out of all microglia, 39% were in contact with plaques, while 61% were away from plaques. Out of plaque-proximal microglia, 83% were LD^+^, whereas only 17% were LD^-^. N=3 5xFAD mice. **c.** In the subiculum of 5xFAD mice, microglia (IBA1, green) exhibited larger cell bodies, shorter processes, and higher LD content (LipidTox, red) when in close proximity to Aβ plaques (MO4, blue), compared to plaque-distant microglia. **d.** Quantification showed a significantly higher morphological index in plaque-proximal compared to plaque-distant microglia in the 5xFAD subiculum; \*\*\**P=* 0.0003. Data represent mean ± SD. Unpaired t-test, N=3 5xFAD mice. **e.** Immunofluorescence for lipid droplets (PLIN2), amyloid plaques (AmyloGlo), and microglia (IBA1) revealed larger LD volume in plaque-proximal microglia in the hippocampus of AD patients compared to plaque-distant microglia. **f.** The average number of IBA1^+^ microglial fragments containing LDs per AD patient, was significantly increased within 10μm from the closest amyloid plaque compared to LD^+^ microglial fragments detected 10-20μm (\*\**P*= 0.003095), 20-30μm (\*\*\**P*= 0.000455), 30-40μm (*P*= 0.000095) or >40µm (\*\**P*= 0.008647) from the closest amyloid plaque. **g.** The sum volume of all LDs within LD^+^ microglial fragments was larger in cells located within 10µm from the closest amyloid plaque compared to LD^+^ microglial fragments detected 10-20μm (\*\**P*= 0.001158), 20-30μm (\*\*\**P*= 0.000241), 30-40μm (\*\*\*\**P*= 0.000065) or >40µm (\*\**P*= 0.001545) from the closest amyloid plaque. For **f-g**: Data represent mean ± SEM. One-way ANOVA with Tukey’s multiple comparison tests, N=6 AD cases (3 males and 3 females). Individual values shown were averaged from 4 z-stacks imaged per patient. **h.** Experimental design for determining the phagocytic capacity and LD load of microglia from 5xFAD and WT female mice (5-7 months old). Microglia were isolated from mouse brains, acutely seeded onto the culture plates for 1 hour, treated with the Aβ^pH^ probe for 30 mins, and with the LipidTox dye for another 30 mins before flow cytometry analysis. **i.** Quantification of LDs in Aβ^pH^- (blue) or vehicle-treated (cyan) microglia from WT mice with fluorescence minus one (FMO) Aβ^pH^ only control (grey). Aβ^pH^ treatment induced an increase in LDs in WT microglia; *\*\*\*P*= 0.0002. **j.** Quantification of LDs in Aβ^pH^- (red) or vehicle-treated (pink) microglia from 5xFAD mice with FMO Aβ^pH^ only control (charcoal). Aβ^pH^ treatment did not induce an increase in LDs in 5xFAD microglia. **k.** Representative dot plots showing LD and Aβ^pH^ uptake by microglia from WT and 5xFAD mice. Microglia from 5xFAD mice showed reduced Aβ^pH^ uptake compared to microglia from WT mice. **l.** Quantification of Aβ^pH^ uptake showed a phagocytic deficit in LD^+^ microglia from 5xFAD compared to LD^+^ microglia from WT mice; \*\**P*= 0.0019. For **i, j** and **l**: Data represent mean ± SD. Unpaired t-tests, cells were pooled from 3 mice per group (3 WT and 3 5xFAD mice) for each of the N=3 experiments.

Plaque-associated microglia have been previously shown to exhibit morphological and molecular differences compared to non-plaque-associated cells in AD^40,41,42^. We specifically asked if direct exposure to Aβ leads to LD formation and changes to phagocytic function in microglia. Microglia isolated from 5-7-month-old female 5xFAD and WT mice were acutely seeded (1 hour) and treated with Aβ^pH^—a pH-dependent fluorescent probe that emits green fluorescence in the acidic lysosomes upon phagocytosis^43^; LDs were then stained with LipidTox, and all cells were analyzed by flow cytometry (**Fig. 2h, Fig. S9a**). Microglia isolated from WT mice showed a significant increase in LD content (4.5-fold) upon direct exposure to Aβ^pH^ (**Fig. 2i**). However, Aβ^pH^ treatment did not cause a significant increase in LDs in microglia isolated from 5xFAD brains (**Fig. 2j**). This could indicate that since 5xFAD mice progressively develop Aβ plaques starting at 2 months of age, chronic microglial exposure to Aβ possibly alters their functional abilities and overall state by the age of 5-7 months, when amyloid deposition is extensive throughout the brain.

Microglial phagocytosis of Aβ is a critical clearance mechanism, and alterations of microglial phagocytic capacity have been reported in chronic inflammation and AD mouse models^15,16^. We therefore investigated if the increase in microglial LDs affects their phagocytic capacity. Live microglia from 5xFAD brains showed a significant (40%) reduction in Aβ^pH^ phagocytosis compared to cells from WT brains (**Fig. S9b-c**). Specifically, out of all microglia, 63.55% and 47.92% were Aβ^pH^**^+^** in WT and 5xFAD, respectively (**Fig. 2k**). Interestingly, LD**^+^** 5xFAD microglia showed impaired phagocytosis compared to LD^+^ WT cells (**Fig. 2l**). Surprisingly, WT microglia showed an increase in LDs due to acute Aβ^pH^ but did not exhibit reduced phagocytic capacity. In conclusion, these experiments demonstrate that LD-laden microglia that are chronically exposed to Aβ exhibit a dysfunctional phagocytic phenotype.

### Direct Aβ exposure is sufficient to significantly alter the microglial lipidome towards lipid droplet formation, independently of inflammatory factors

We were surprised to discover that in microglia from 5-7-month-old WT brains, even acute direct exposure to Aβ *ex vivo* was sufficient to induce large LD formation. Thus, we asked if direct Aβ exposure is sufficient to promote LD formation and/or regulate broad metabolic changes in microglia independently of other inflammatory factors. We isolated CD11b^+^ primary microglia from 5-7-month-old WT mice, cultured them *in vitro* for 7-10 days and treated them with Aβ aggregates for 1, 12, and 24 hours before assaying both cellular and secretory lipids and metabolites by mass spectrometry-based MRM-profiling (**Fig. 3a**). We again screened over 1370 lipid species categorized into 10 main classes including FFAs, ceramides, and TAGs **(Supplementary Tables ST6, ST7**) as well as over 700 metabolites (**Supplementary Tables ST8, S9**). We found considerable differences in the microglial lipidome between Aβ- and vehicle-treated cells, at 1 hour and 24 hours of Aβ treatment, respectively, suggesting a distinct lipidomic transition occurring within microglia following exposure to Aβ (**Fig. 3b**).

**Fig. 3:**
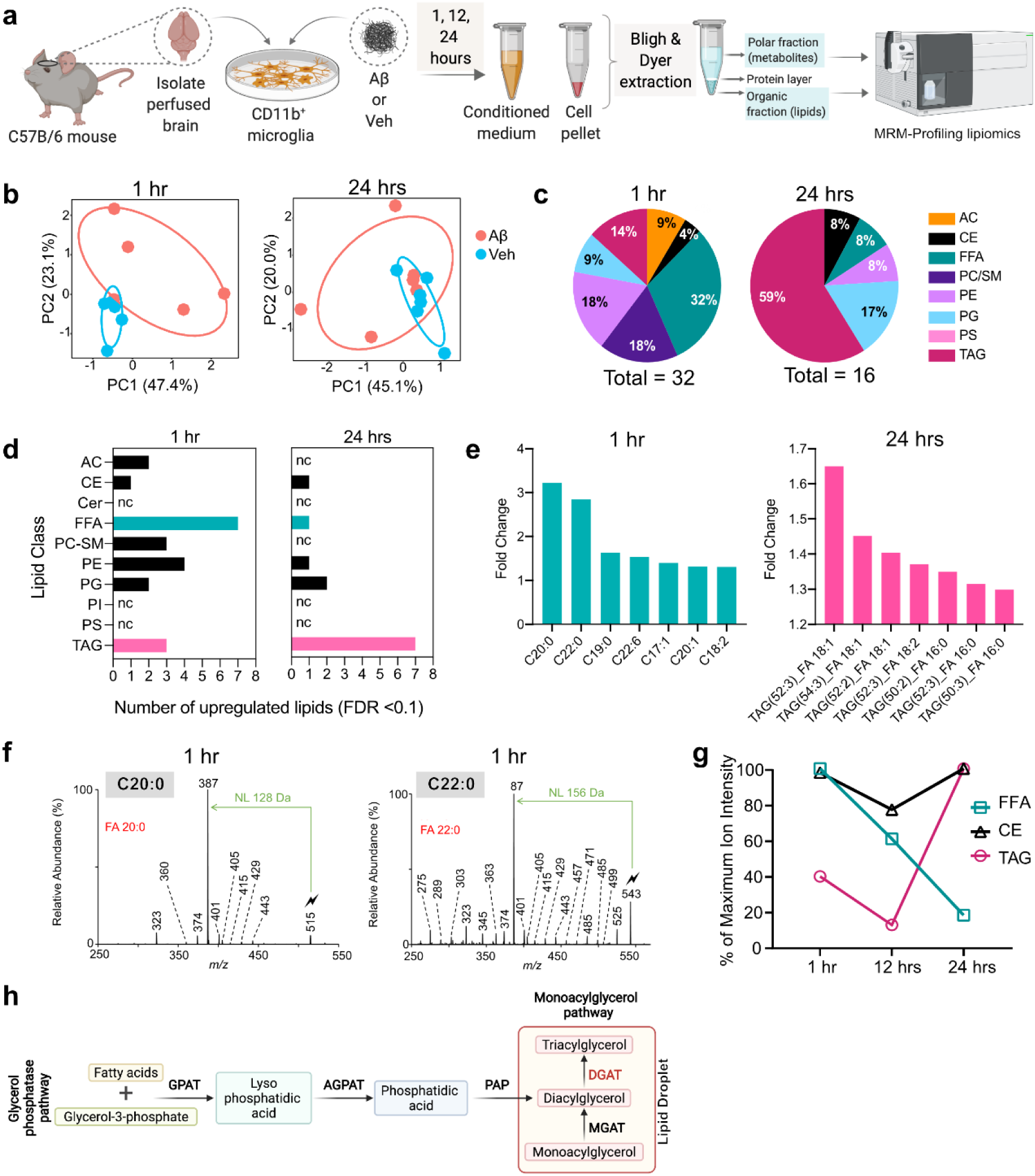
Aβ induces profound changes to the microglial lipidome and metabolome *in vitro*, resulting in LD formation. **a.** Experimental design for the global lipidomic profiling experiment performed on Aβ- and vehicle-treated primary mouse microglia. Cells were isolated from ∼7-month-old C56BL/6J perfused mouse brains and cultured in growth medium containing TGF-β, IL-34, and cholesterol. Cells isolated from each brain were split and treated with 500 nM Aβ or vehicle for 1, 12, and 24 hours, followed by lipid and metabolite extraction from conditioned media and cell pellets, which were run on the Agilent triple quadrupole mass spectrometer. Lipids and metabolites were identified in the samples using MRM-profiling. Each experiment was repeated 5 or 6 times resulting in: N=5 mice were used for 1-hour treatments, N=6 mice for 12 hours, and N=6 mice for 24 hours treatments. **b.** PCA demonstrating the variation in microglial lipidomes both within and between groups (Aβ or vehicle treated microglia) at 1 and 24 hours of treatment. **c.** The distribution of significantly different lipid classes identified in microglia at 1 and 24 hours of Aβ treatment, compared to vehicle treated cells. 32% of the differentially regulated lipids at 1 hour were FFA, whereas 59% of the differentially regulated lipids at 24 hours were TAGs. **d.** Upregulated lipid classes at 1 and 24 hours of Aβ treatment compared to vehicle, showed FFAs and TAGs were the most abundant lipids, respectively. **e.** Individual lipid species belonging to FFAs and TAGs that were upregulated at 1 and 24 hours of Aβ treatment, compared to vehicle. Long-chain saturated FFAs C20:0, C22:0, and C19:0 were the top 3 upregulated FFAs within the first 1-hour of Aβ treatment, while neutral lipids TAG(52:3)_FA 18:1, TAG(54:3)_FA 18:1, and TAG(52:2)_FA 18:1 were the top three upregulated TAGs with prolonged 24-hour Aβ treatment, both compared to vehicle. **f.** Structural identification and confirmation of the C20:0 and C22:0 lipids in the 1-hour Aβ-treated microglial samples, using the gas-phase ion/ion chemistry (see **Supplementary Results**). **g.** Percentage changes of maximum ion intensities as a quantitative measure of changes in the respective amounts of FFAs (green), TAGs (magenta), and CEs (black) in microglial cells at 1, 12, and 24 hours of Aβ treatment. The reduction in FFAs was followed by an increase in TAGs and CEs – major components of LDs – suggesting a gradual conversion of FFAs to TAGs towards LD formation. **h.** Convergent pathways for TAG biosynthesis. Glycerol-3-phosphate acyltransferase (GPAT); acylglycerol-3-phosphate acyltransferase (AGPAT); phosphatidic acid phosphatase (PAP); monoacylglycerol acyltransferase (MGAT); and the final rate-limiting enzyme diacylglycerol acyltransferase (DGAT) that is needed for TAGs production and is involved in LD formation.

While we observed dramatic changes in several lipid classes, FFAs were the most differentially-regulated lipid class with acute (1 hour) Aβ exposure (**Fig. 3c-d**), with very long-chain saturated FFAs, C20:0, C22:0, and C19:0 being the most upregulated lipids at this time point (**Fig. 3e; S10a**). We also used novel gas-phase ion/ion chemistry (**Supplementary Results**) to structurally validate these highly-expressed saturated FFAs and confirmed that these specific saturated FFA structures were indeed directly synthesized in microglia with acute (1 hour) Aβ exposure (**Fig. 3f, S11a-b**). Interestingly, the cells did not maintain their saturated FFA repertoire with prolonged Aβ exposure (24 hours) and transitioned to TAGs, which were the most differentially-regulated lipid class at this time point, with TAG(52:3), TAG(54:3), and TAG(52:2) being the most upregulated species (**Fig. 3c-e; S10b**). In addition, MRM-profiling confirmed the increase of specific TAGs in Aβ-treated microglia at the 24-hour time point compared to vehicle-treated microglia (**Fig. 3e, Supplementary Table ST10**). Furthermore, at 12 hours of Aβ treatment, in addition to FFAs, cholesteryl esters (CEs) were the second most differentially-regulated lipid class, with very long chain CEs 20:2, 24:1, and 16:3 being the top upregulated CEs in microglia (**Fig. S10c-e**). Neutral lipids like TAGs and CEs form the core of LDs and are involved in energy storage and fatty acid metabolism in cells. Importantly, we quantified the total amount of upregulated FFAs, CEs, and TAGs as percent change of maximum ion intensity and verified that the reduction in FFAs from 1 hour to 24 hours was followed by a concomitant increase in TAGs and CEs at 24 hours (**Fig. 3g**). This increase in core LD components suggested that the cells likely activated metabolic pathways towards LD formation. Taken together with acute treatment of Aβ *ex vivo* (**Fig. 2b**), these results suggest a direct effect of Aβ-aggregate exposure to promote LD formation in microglia. In contrast to the cellular lipidome, we did not find many lipids differentially regulated in the microglial conditioned media (secretory lipids) with Aβ treatment - none at 1 hour, five at 12 hours, and one at 24 hours, respectively (**Supplementary Table ST7**). Given that both lipids and metabolites work together to activate cellular metabolic pathways, we also evaluated changes in microglial metabolite profiles with Aβ exposure. Metabolites corresponding to alanine, aspartate, and glutamate metabolism, arginine biosynthesis, etc. were differentially regulated in microglia exposed to Aβ (**Fig. S12, Supplementary Table ST11**). The metabolome of the microglial conditioned media exhibited pronounced differences following 12 and 24 hours of Aβ exposure, as seen in the PCA plots (**Fig. S13a**). Metabolites related to phenylalanine/tyrosine/tryptophan biosynthesis and glycine/serine/threonine biosynthesis pathways were differentially regulated in the microglial conditioned media due to Aβ (**Fig. S13b-c, Supplementary Table ST12**). Taken together, the Aβ-treated cellular and secreted metabolites overlapped with metabolic pathways in 5xFAD compared to WT (**Fig. S2**). This suggests that Aβ plays a direct role in upregulating TAGs, CEs and associated metabolites towards LD formation in AD. TAGs can be synthesized via two major pathways involving the conversion of FFAs to TAGs via several intermediates: 1) the glycerol phosphatase pathway or 2) the monoacylglycerol pathway (**Fig. 3h**)^44^. Since TAGs constitute the major neutral lipid core of LDs and are integral to their structure and function, we hypothesized that these pathways could be directly involved with the dramatic increase in TAGs following Aβ exposure and likely also with LD formation in AD.

### The DGAT2 pathway is required for Aβ-induced lipid droplet formation in AD microglia, and inhibiting it rescues microglial phagocytic impairment and reduced plaque load in AD

The diacylglycerol O-acyltransferase (DGAT) enzymes catalyze the final rate-determining step in the biosynthesis of TAGs. The DGAT2 enzyme is evolutionarily conserved across eukaryotes and has the predominant and ancient function for mediating TAG synthesis from fatty acids^45^. In addition, DGAT2 localizes around LDs, where it is required for LD-localized TAG synthesis^46,47,48^. Thus, we investigated whether DGAT2 is involved in the formation of Aβ-induced LDs in microglia (**Fig. 4a**). We immunostained 5xFAD and WT brain tissues for DGAT2, LDs, Aβ plaques, and microglia. We found that LD-laden microglia surrounding the Aβ plaques expressed DGAT2 in 5xFAD tissue (**Fig. 4b**), despite *Dgat2* mRNA being downregulated in 5xFAD microglia (**Fig. S14a**), we detected abundant DGAT2^+^ microglia in the subiculum region of the 5xFAD brain (**Fig. S14b, Fig. 4c**). We also immunostained human AD and NS hippocampal tissue for Aβ plaques, LD, DGAT2, and microglia. We found that plaque-associated microglia with increased LD content exhibited higher DGAT2 levels in AD brains compared to microglia from NS brains (**Fig. 4d**).

**Fig. 4:**
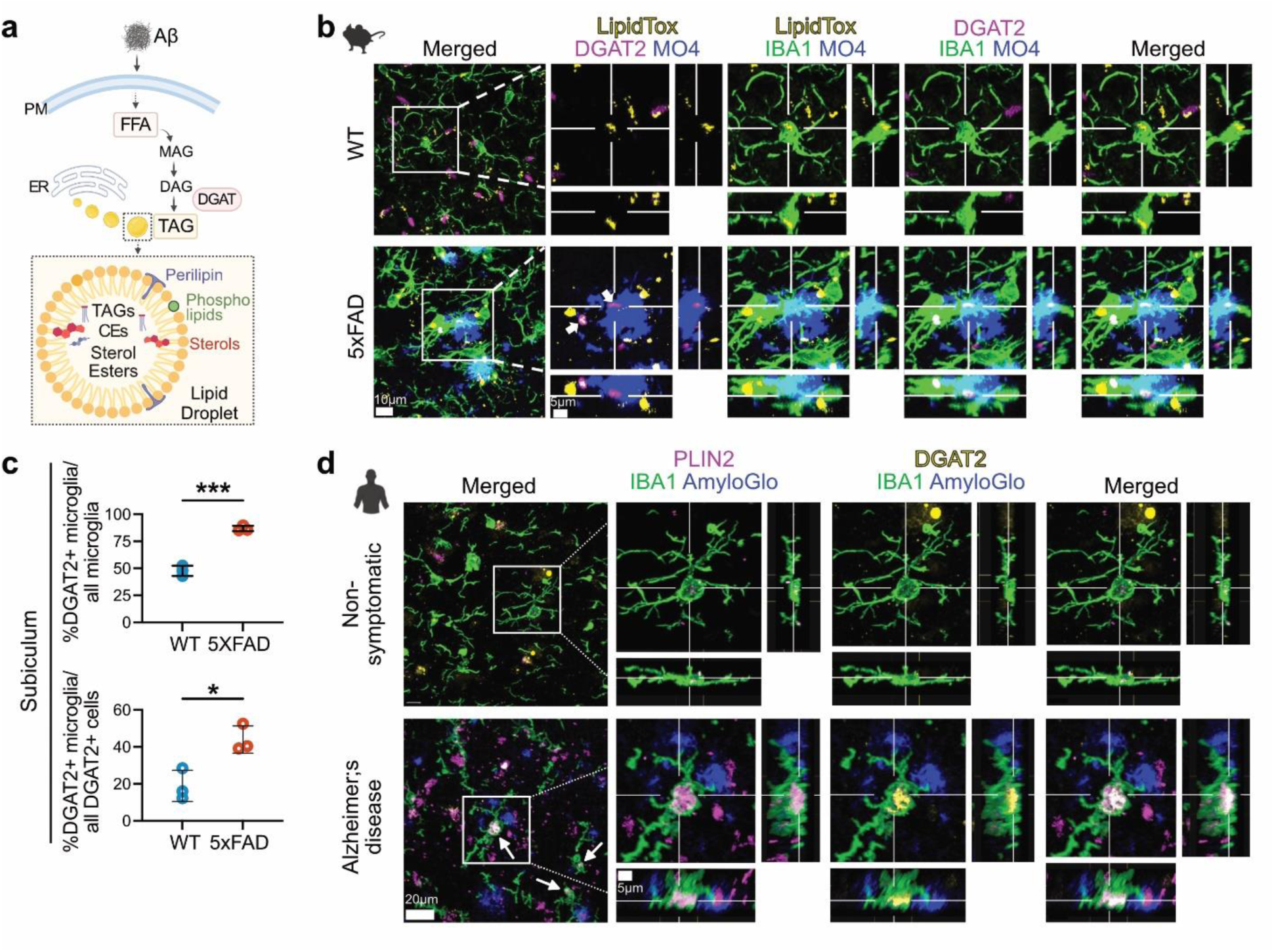
DGAT2 enzyme is highly expressed in LD-laden plaque proximal microglia in mouse and human AD brain. **a.** Proposed mechanism for Aβ-induced LD formation in microglia. Microglial exposure to Aβ induces an upregulation of FFAs that are converted to TAGs within LDs via the DGAT2 pathway. **b.** Immunofluorescence staining of microglia (IBA1), LDs (LipidTox), DGAT2, and Aβ plaques (MO4) in the hippocampal subicular region of 5xFAD and WT mouse brains. Increased DGAT2 is shown in microglia associated with amyloid plaques. **c.** Quantification showing a significant increase in % of DGAT2^+^ microglia out of all microglia and out of all DGAT2^+^ cells in the mouse subiculum in the 5xFAD tissue vs. WT; \**P=* 0.0181, *\*\*\*P=* 0.0002. Data represent mean ± SD. Unpaired t-test, N=3 mice per group. **d.** DGAT2 expression in LD^+^ microglia in close proximity to amyloid plaques in hippocampal FFPE tissue from human AD and NS cases (N=4 per group). Immunofluorescence was performed on 15µm-thick human hippocampal sections for the detection of DGAT2, lipid droplets (PLIN2), amyloid plaques (AmyloGlo) and microglia (IBA1). Increased DGAT2 signal (yellow) was detected in plaque-proximal LD^+^ microglia in AD cases (arrows), compared to NS controls. Cross-sections of the selected microglial cells in the white boxes demonstrate representative example of increased DGAT2 signal in close proximity to a large PLIN2-labeled LD inside a plaque-proximal microglial cell in AD, and of a cell from a non-symptomatic case.

Further, we used a DGAT2 inhibitor (D2i) to determine if reducing DGAT2 function affects microglial LD content and Aβ-specific phagocytosis. Microglia from 5-7-month-old female 5xFAD and WT mice were acutely seeded (1 hour) and treated with Aβ^pH^ and LipidTox sequentially, in the presence of D2i or vehicle, after which cells were analyzed by flow cytometry (**Fig. 5a**). Both WT and 5xFAD microglia showed a significant decrease in LDs upon D2i treatment *in vitro* (approx. 51% and 57% decrease, respectively, **Fig. 5b**). Further, there was a significant reduction in Aβ-induced LDs with D2i treatment in WT microglia, thereby confirming that Aβ-induced LD formation requires the DGAT2 pathway (**Fig. 5c-d, Fig. S15a**). Interestingly, we did not observe any significant differences in Aβ-induced LDs with D2i treatment in 5xFAD microglia (**Fig. 5c-d, Fig. S15b)**, similar to our results showing that Aβ^pH^ treatment did not affect LD load in microglia isolated from 5xFAD brains. Together, these results suggest an underlying Aβ-induced LD saturation mechanism that allows DGAT2-mediated microglial LD formation in 5xFAD brain after chronic amyloid exposure *in vivo*, but prevents further LD accumulation upon additional *in vitro* Aβ exposure.

**Fig. 5:**
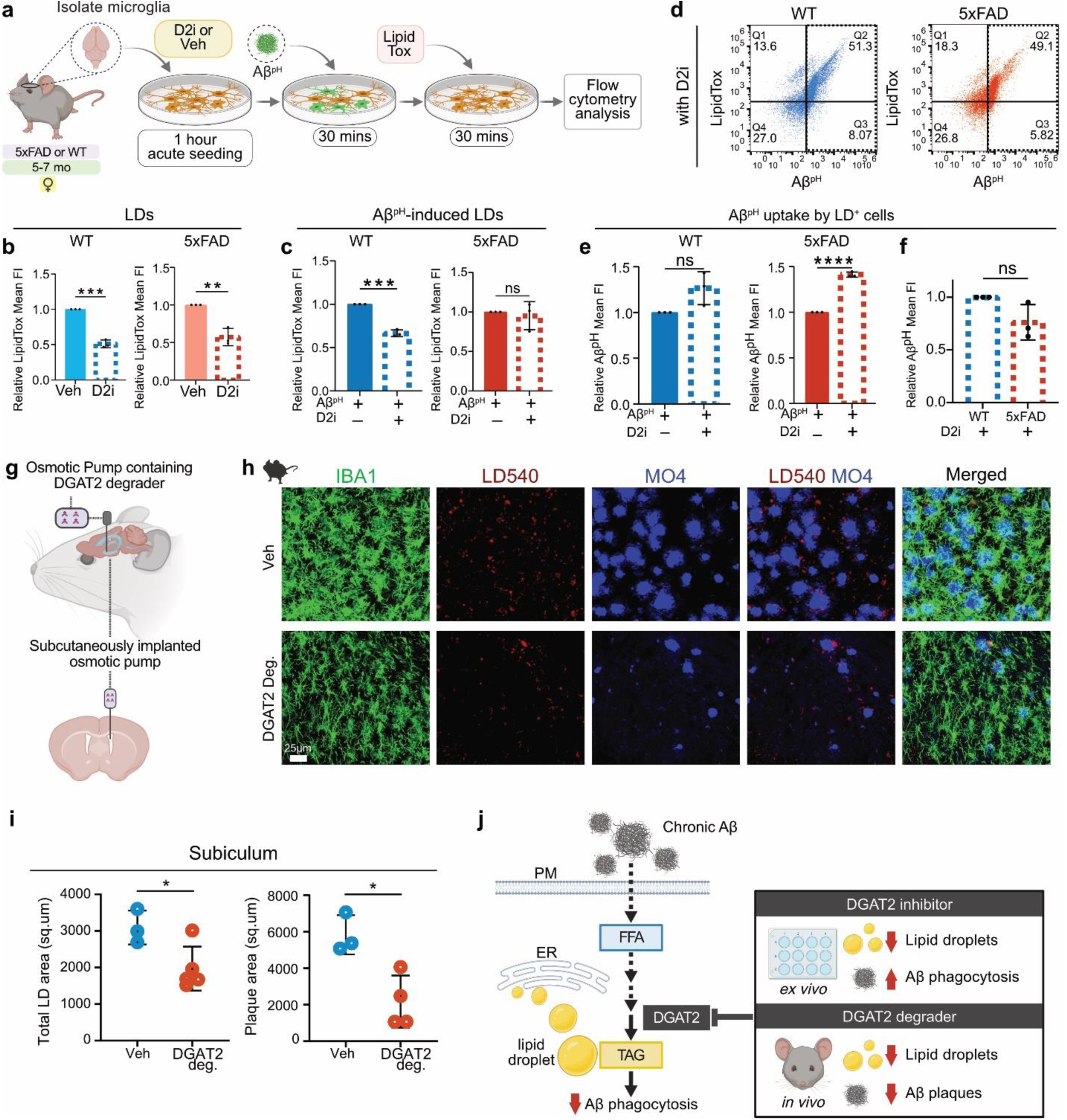
DGAT2 enzyme is required for Aβ-induced LD formation, and inhibiting or degrading it restores Aβ phagocytosis, decreases LD and plaque burden in microglia. **a.** Experimental design for determining the phagocytic capacity and LD load of microglia from 5xFAD and WT female mice (5-7 months old). Microglia were isolated from mouse brains, acutely seeded onto the culture plates with D2i or Veh for 1 hour, followed by sequential treatment with Aβ^pH^ probe and LipidTox dye, in presence of D2i or Veh, before flow cytometry analysis. **b.** DGAT2 inhibitor (D2i) treatment reduced LDs in cultured microglia from WT and 5xFAD brains; \*\*\**P=* 0.0001, *\*\*P=* 0.0029. **c.** Quantification showed that D2i treatment reduced LD formation upon Aβ exposure in microglia from WT mice but not in cells from 5xFAD mice; \*\*\**P*= 0.0001. **d.** Representative dot plots showing LD and Aβ^pH^ uptake by microglia treated with D2i from WT and 5xFAD mice. **e.** LD^+^ microglia from WT mice showed a slight but non-significant increase in Aβ^pH^ uptake with D2i, while LD+ microglia from 5xFAD mice showed a significant increase in Aβ^pH^ uptake with D2i; \*\*\*\**P*= 0.000014. **f.** Direct comparison of the effect of D2i treatment on Aβ^pH^ uptake by LD^+^ microglia from WT and 5xFAD showed that inhibiting DGAT2 restored the phagocytic performance of 5xFAD microglia, making it comparable to that of WT cells. For **c**, **d, e,** and **f**: Data represent mean ± SD. Unpaired t-tests, cells were pooled from 3 mice per group (3 WT and 3 5xFAD mice) for each of the N=3 experiments. **g.** Schematic showing the delivery of DGAT2 degrader into the lateral ventricles of 18-24 months old 5xFAD mice using subcutaneously implanted osmotic pumps. **h.** Immunofluorescence for microglia (IBA1), LDs (LD540), Aβ plaques (MO4) in the hippocampal subicular region of vehicle and DGAT2 degrader-treated 5xFAD mice showed evident LDs and Aβ plaques reduction following DGAT2 degrader treatment. **i.** Quantification showed a significant decrease in total LD area and Aβ plaque area in the subiculum of 5xFAD mice treated with DGAT2 degrader vs. vehicle; \**P<*0.05. Data represent mean ± SD. Unpaired t-test, N=5 mice received DGAT2 degrader and N=3 mice received vehicle treatment. **j.** Model proposing DGAT2 as the target in AD for Aβ-induced LD formation and phagocytic dysfunction in microglia. Inhibition or degradation of DGAT2 resulted in increased Aβ uptake and reduced plaque burden, respectively, while reducing LD load.

Next, we examined if D2i affected microglial phagocytic capacity. LD^+^ microglia from WT brains showed a slight but non-significant increase in Aβ^pH^ uptake with D2i compared to vehicle-treated cells (**Fig. 5e; Fig. S15c**). In contrast, LD^+^ 5xFAD microglia showed a significant increase (1.41-fold) in Aβ^pH^ uptake with D2i, which was similar to WT microglia in the presence of the inhibitor (**Fig. 5e-f; Fig. S15f).** Furthermore, the overall phagocytic capacity of 5xFAD microglia was improved with D2i treatment (**Fig. S15c**) compared to vehicle-treated cells, although again there was no significant difference between WT and 5xFAD microglia in the presence of the inhibitor (**Fig. S15d-e**). Thus, the phagocytic dysfunction of LD-laden 5xFAD microglia was attenuated in the presence of D2i.

To investigate how targeting DGAT2 *in vivo* might affect amyloid pathology, we developed a degrader of DGAT2^49^ and continuously infused it into the lateral ventricles of 18-24 month old 5xFAD mice (**Fig. 5g**). Animals that received the DGAT2 degrader over a period of 1 week showed a significant reduction in total LD area [36%] in the subicular hippocampal region compared to the age-matched vehicle-treated control animals (**Fig. 5h,i, Fig. S16a**). Strikingly, the DGAT2 degrader also drastically reduced the amyloid load in the 5xFAD mice by 63% compared to mice that received the vehicle treatment (**Fig. 5i, Fig. S16b**). Taken together, these findings indicate that degrading DGAT2 profoundly reduces LD content and improves amyloid pathology even in chronically inflamed 18-24 month old 5xFAD mice. which typically presents itself with an excessive amount of brain amyloid deposition and advanced disease.

In summary, we have discovered a novel Aβ-mediated mechanism that promotes LD formation in microglia and identified the DGAT2 pathway as a novel target for therapeutic intervention to reduce phagocytic impairment of microglia and promote amyloid clearance or limit amyloid accumulation in the AD brain (**Fig. 5j**).

## DISCUSSION

Over the past two decades, a plethora of studies have revealed important microglial contributions to practically every neurological disorder or disease, with roles ranging from protective to harmful^50^. This collective body of work, together with an increasing number of microglial transcriptomic profiling studies, have underscored their multidimensional nature and their impressive capacity to react to any signal presented to them in an age–, sex–, disease stage–, and overall context–specific manner^50^. In this study, we showed that in the context of AD pathology, chronic exposure to Aβ promotes specific metabolic changes in microglia that make them progressively accumulate LDs and render them incapable of contributing to Aβ clearance. LD-laden microglia increase in numbers in an age- (5-7-month old), sex- (female), and brain region-specific manner (subiculum) in 5xFAD mice. These microglia are closely associated with Aβ plaques and have a unique morphology comprised of larger cell bodies and shorter processes in both 5xFAD and human AD brains. Importantly, LD-laden microglia exhibit reduced phagocytosis of Aβ, a critical functional consequence of LD accumulation. Further, by challenging primary WT microglia directly with Aβ, we discovered that even acute exposure to Aβ was sufficient to shift their lipidomic composition towards reduced FFA and increased TAG content – a major component of LDs. We thus identified DGAT2, a key enzyme for the conversion of FFAs to TAGs, as an essential mediator of LD formation in microglia, and showed its increased abundance in 5xFAD as well as human AD brain tissue. Crucially, pharmacological inhibition of DGAT2 improved Aβ phagocytosis *ex vivo*, and DGAT2 degradation drastically reduced plaque load *in vivo*, thus unraveling a new molecular target and mechanism related to microglial dysfunction in AD.

Recent studies focused on transcriptomic changes occurring in microglia at the single-cell level due to Aβ. While such studies demonstrate the extent of glial heterogeneity in a specific environment based upon highly-abundant and actively-transcribed genes in each subpopulation, they do not fully capture the functional aspects of these cell populations, which are directly determined by bioactive molecules such as proteins, lipids, and other metabolites. We performed the first-ever large-scale lipidomic and metabolomic characterization of microglia with acute exposure to Aβ *in vitro* and in chronic 5xFAD mice, profiling over 1300 lipid species and 700 metabolites. Our data, which will be made available online (http://microgliaomics-chopralab.appspot.com), showed an increase in long-chain saturated FFAs (specifically C20:0, C19:0, and C22:0) upon 1h Aβ exposure; however, longer treatment with Aβ (24h) resulted in an abundance of long-chain TAGs. Previous studies showed that saturated FFAs are extremely cytotoxic to cells^51^. One way that cells overcome FFA-induced cytotoxicity is by metabolizing them into TAGs^52^, which are integral components of the LD core. Microglia are thus able to circumvent lipid-mediated cytotoxicity via the production and accumulation of LDs. Interestingly, we recently found that reactive astrocytes secrete long-chain saturated FFAs that kill injured neurons and oligodendrocytes *in vitro* and *in vivo*^37^. While injured neurons and oligodendrocytes typically have a low capacity for consuming FFAs and sequestering them into LDs^53^, making them more vulnerable to lipid-induced cytotoxicity in chronic inflammation^54,55^, our findings demonstrate that microglia can metabolize these toxic molecules into TAGs. This response could be one way that microglia protect themselves and neurons from the cytotoxic environment of the chronically-inflamed AD brain; however, it seems to affect their own morphology and functional state as well.

Changes in microglial morphology have been described in numerous inflammatory, degenerative, and injury paradigms as indications of changes in their overall state and reactivity. In our study, LD-laden microglia were closely associated with Aβ plaques and exhibited shorter processes and larger cell bodies in both 5xFAD mouse and human AD brains. These unique LD-laden microglia were primarily located in the hippocampal subicular regions of 5xFAD mice and in postmortem human hippocampi from AD patients, which are the sites of the earliest atrophy seen during AD pathogenesis^56^. In addition to the protective roles of some lipids^57^, their accumulation is also linked to cellular senescence^58^. For example, in an obesity mouse model, senescent glia were found to accumulate LDs, which contributed to impaired neurogenesis^59^. Whether these LD-laden microglia are ultimately detrimental or protective to the tissue due to their dysfunctional, hyper-reactive, and/or senescent state in the context of chronic inflammation remains elusive and may well depend upon the extent or nature of metabolic changes that microglia experience in the context of different challenges. Overall, these observations open new avenues for exploring the functional roles of plaque-associated LD-laden microglia and their involvement in AD progression, as well as the functional significance of LD accumulation in microglia in other neuropathologies.

Phagocytosis is a major innate immune mechanism contributing to Αβ clearance, and its failure favors Αβ accumulation in the brain. Late-onset AD, which accounts for ∼95% of AD cases, is associated with impaired Αβ clearance^60^. Microglia directly engulf and degrade Aβ^61,62^, but their capacity for efficient phagocytosis declines with aging, chronic exposure to Αβ, and during sustained inflammation in the AD brain^63,16,17^. Indeed, acute exposure of primary microglia to Aβ increased their inflammatory cytokine secretion and phagocytic performance, while chronic exposure reduced phagocytosis and induced immune tolerance as well as a metabolic shift from oxidative phosphorylation to glycolysis^64^. Interestingly, restoration of microglial phagocytosis improved cognitive function in aged mice and promoted a homeostatic microglial transcriptional signature^65^.

Plaque–associated microglia have direct access to remove aggregated amyloid through phagocytosis. Prior studies have proposed that these cells are compacting amyloid plaques through phagocytic receptors such as TREM2^14^, (a receptor expressed on myeloid cells, for which gene variants have been associated to increased AD risk)^14,8^, Axl, and Mer^66^. Genetic deletion of any of these receptors in mouse models of AD resulted in less compact/more diffuse plaques, and was associated with increased neuritic dystrophy around Αβ deposits^14,66^. These studies suggested that microglia build a neuroprotective barrier that limits plaque growth through their phagocytic receptors, and further proposed microglia as plaque “compactors” that uptake Αβ and condense it in their lysosomes to eventually re-deposit it as dense core plaques^66^. We found that plaque– associated microglia progressively accumulated more LDs, and that cells with increased LD content were less capable of Aβ phagocytosis. LD-laden microglia found in the aged brain also demonstrated phagocytic deficits^20^, but the molecular mechanisms linking the appearance of these organelles with functional deficits in microglia were unclear. We discovered that Aβ alone can shift the metabolic equilibrium within microglia to convert FFAs to TAGs, one of the main components of LDs. To decipher if this catalytic conversion that is known to contribute to LD formation^52,67^ is also responsible for microglial dysfunction, we blocked DGAT2, a key enzyme for this pathway. Although we found the DGAT2 mRNA significantly decreased in 5xFAD microglia, the protein was abundant in plaque-associated LD-laden microglia in both mouse and human AD brains, underscoring how relying solely on transcript levels might mask important ongoing cellular functions. Importantly, blocking DGAT2 was sufficient to restore phagocytosis in 5xFAD microglia, suggesting that phagocytic dysfunction is the consequence, rather than the cause, of chronic LD accumulation. In support of our findings, a study in macrophages showed that their phagocytic function is also dependent upon the availability of FFAs that are generated upon degradation of LDs, suggesting that LD accumulation somehow limits the phagocytic capacity of all tissue-resident macrophages^68^. Furthermore, using a complementary DGAT2 protein degrader approach also reduced LD load *in vivo* in 18-24 month 5xFAD mice, underscoring the importance of this molecular target for LD accumulation. Crucially, its very rapid (only 1 week of treatment) and profound reduction (63%) of amyloid load in older animals highlights DGAT2 as a promising target to restore microglia-mediated Αβ clearance and decrease amyloid deposition even in AD patients of advanced age and/or with increased amyloid pathology.

Delineating the molecular mechanisms that drive LD accumulation and their downstream impact on cellular functions and overall tissue health is paramount for the design of novel therapeutic strategies for AD and other neurodegenerative diseases. Here, we have identified one such druggable molecule that regulates TAG formation and LD accumulation in microglia while also directly impacting their function. Unlike other acyltransferases, such as GPAT and AGPAT, that are upstream and involved in the glycerol phosphatase pathway, the activity of DGAT is the direct rate-limiting step for the biosynthesis of TAGs^44^. Therefore, since disrupting DGAT2 alone was sufficient to improve microglial phagocytosis of Aβ, and reduce brain amyloid load, targeting DGAT2 emerges as a prime target for regulating the phagocytic activity of LD-laden microglia compared to other candidate enzymes. Our study thus provides the first proof-of-principle that disrupting DGAT2 can be a highly effective strategy for promoting the protective role of microglia in AD, and possibly in other neurodegenerative diseases with excessive protein aggregation and LD deposition.

## DATA AVAILABILITY

Supplemental tables, figures, and associated content is available with the manuscript. All data analysis is available on GitHub (https://github.com/chopralab/microglia_omics). A web application has been developed for exploring lipid and metabolite mass spectrometry data that will be publicly available at http://microgliaomics-chopralab.appspot.com (for review, username: admin, password: Review). The accession information for raw lipid and metabolite mass spectrometry data is MassIVE MSV000089458: https://massive.ucsd.edu/ProteoSAFe/dataset.jsp?task=0f7bd7cfaf504869bfac786e4184105e.

## CODE AVAILIBILITY

All of the analysis codes for the lipidomics and metabolomics experiments are available on Github at https://github.com/chopralab/microglia_omics.

## ACKNOWLEDGEMENTS

We thank the following individuals for their input and assistance: Ms. Anisa Dunham for help with animal breeding and maintenance, as well as PCR experiments; Dr. Christina Ferreira at the Purdue Metabolomics Facility for assistance with lipid mass spectrometry; Dr. Scott McLuckey for access to a modified Sciex QTRAP 4000 triple quadrupole/linear ion trap mass spectrometer to perform ion/ion reactions; Dr. Shane Tichy for his support, and Agilent Technologies for their gift of the Triple Quadrupole LC/MS to the Chopra Laboratory; Dr. J. Paul Robinson and Ms. Kathy Ragheb at the Purdue University Cytometry Laboratories for flow cytometry services; Dr. Chris Nelson in the Department of Neurosciences of the Lerner Research Institute of Cleveland Clinic for editorial assistance. This work was supported by the United States Department of Defense USAMRAA award W81XWH2010665 through the Peer Reviewed Alzheimer’s Research Program, NIH National Institute of Mental Health award R01MH128866 and National Center for Advancing Translational Sciences ASPIRE awards to G.C.; the NIH National Institute for Neurological Disorders and Stroke award R01NS112526, and NIH National Institute on Alcohol Abuse and Alcoholism award P50AA024333 to D.D. Additional support, in part, by the Stark Neurosciences Research Institute, the Indiana Alzheimer Disease Center, Eli Lilly and Company; the Indiana Clinical and Translational Sciences Institute grant UL1TR002529 from the NIH, National Center for Advancing Translational Sciences. We also acknowledge the Pathology Research Core in the Robert J. Tomsich Pathology and Laboratory Medicine Institute of Cleveland Clinic for their human tissue services; the Clinical Core of Cleveland Clinic’s Northern Ohio Alcohol Center funded by NIH grant P50AA024333; the Purdue University Center for Cancer Research funded by NIH grant P30 CA023168. We also thank current and prior members of the Davalos and Chopra laboratories for critical discussions and day-to-day assistance with experiments included in this manuscript. The content is solely the responsibility of the authors and does not necessarily represent the official views of the National Institutes of Health. Select illustrations in figures were made using BioRender.

## DECLARATION OF INTERESTS

G.C. is the Director of the Merck-Purdue Center for Measurement Science funded by Merck Sharp & Dohme, a subsidiary of Merck and a co-founder of Meditati Inc., a startup developing smart drugs for mental health indications. The remaining authors declare no competing interests.

## SUPPLEMENTARY TABLE LEGENDS

**Supplementary Table ST1: Analyzed lipidomics data of 5xFAD vs. WT microglia, including female and male samples.** Tables show the differential lipid profiles in 5xFAD vs. WT microglia samples, organized by most to least differentially-expressed lipids. Samples N1, N2, N3 belong to female mice and samples N4, N5 belong to male mice. P and FDR values are also provided and lipids with FDR<0.1 were considered to be significant.

**Supplementary Table ST2: Analyzed lipidomics data of 5xFAD vs. WT microglia, female only.** Table shows the differential lipid in 5xFAD vs. WT female microglia samples, organized by most to least differentially-expressed lipids. P and FDR values are also provided and lipids with FDR<0.1 were considered to be significant.

**Supplementary Table ST3: Raw MRM lipidomics data of 5xFAD vs. WT microglia.** Table lists the MRM transitions screened for TAGs and the respective ion intensity values, organized by MRM transition.

**Supplementary Table ST4: Analyzed metabolomics data for 5xFAD vs. WT microglia (male and female).** Tables show the differential metabolites in 5xFAD vs. WT microglia, organized by most to least differentially-expressed metabolites. P and FDR values are also provided and metabolites with FDR<0.1 were considered to be significant.

**Supplementary Table ST5: Pathway analysis for 5xFAD vs. WT microglial metabolites.** A list of all pathways matched to the differentially-expressed metabolites (FDR<0.1) is provided in the table. The pathways with P<0.05 were considered to be significant and are highlighted in the scatter plots in the supplementary figures.

**Supplementary Table ST6: Analyzed lipidomics data for Aβ- vs. vehicle-treated primary microglial cells at 1, 12, and 24 hours.** Tables show the differential lipid profiles in Aβ- vs. vehicle-treated cells at the three different time points, organized by most to least differentially-expressed lipids. P and FDR values are also provided and lipids with FDR<0.1 were considered to be significant.

**Supplementary Table ST7: Analyzed lipidomics data of Aβ- vs. vehicle-treated primary microglial cell conditioned media at 1, 12, and 24 hours.** Tables show the differential lipid profiles in Aβ- vs. vehicle-treated conditioned media samples at the three different time points, organized by most to least differentially-expressed lipids. P and FDR values are also provided and lipids with FDR<0.1 were considered to be significant.

**Supplementary Table ST8: Analyzed metabolomics data of Aβ- vs. vehicle-treated primary microglial cells at 1, 12, and 24 hours.** Tables show the differential metabolites in Aβ- vs. vehicle-treated cells at the three different time points, organized by most to least differentially-expressed metabolites. P and FDR values are also provided and metabolites with FDR<0.1 were considered to be significant.

**Supplementary Table ST9: Analyzed metabolomics data of Aβ- vs. vehicle-treated primary microglial cell conditioned media at 1, 12, and 24 hours.** Tables show the differential metabolites in Aβ- vs. vehicle-treated conditioned media samples at the three different time points, organized by most to least differentially-expressed metabolites. P and FDR values are also provided and metabolites with FDR<0.1 were considered to be significant.

**Supplementary Table ST10: Raw MRM lipidomics data of Aβ- vs. vehicle-treated primary microglial cell conditioned media at 1 and 24 hours.** The table lists the MRM transitions screened for TAGs and the respective ion intensity values at two different time points, organized by MRM transition.

**Supplementary Table ST11: Pathway analysis for cell metabolites at 1, 12, and 24 hours of Aβ- vs. vehicle-treated primary microglia cultures.** List of all pathways matched to the differentially-expressed metabolites (FDR<0.1) are highlighted in the tables. The pathways with P<0.05 were considered to be significant and are highlighted in the scatter plots in the supplementary figures.

**Supplementary Table ST12: Pathway analysis for media metabolites at 1, 12, and 24 hours of Aβ- vs. vehicle-treated primary microglia cultures.** List of all pathways matched to the differentially-expressed metabolites (FDR<0.1) are highlighted in the tables. The pathways with P<0.05 were considered to be significant and are highlighted in the scatter plots in the supplementary figures.

## SUPPLEMENTARY MOVIE LEGEND

**Supplementary Movie 1: 3D rendered plaque-proximal microglia show increased LDs in human AD patients.** Representative confocal z-stack for which 3D “Surfaces” were made in Imaris for IBA1, PLIN2 and AmyloGlo channels. Only LDs in microglia (red) are shown. Plaque-proximal microglia (yellow) defined by a 0-10μm distance from the closest plaque (purple) contain increased LDs (red) compared to plaque-distant (>10μm from closest plaque) LD+ microglia (green). Plaque-proximal LD-microglia are also shown in orange.

## STAR Methods

### Animals

C57BL/6J and 5xFAD mice were obtained from the Jackson Laboratory and were maintained in a pathogen free facility. All experiments involving mice were performed in accordance with the Purdue University’s Institutional Animal Care and Use Committee (IACUC) guidelines.

### Primary mouse microglia isolation and culture from adult mouse brains

Primary microglia from adult mouse brains were isolated and cultured per a previously-described protocol^43^. Briefly, CD11b^+^ primary microglia were isolated from adult mice (both male and female) and cultured as described previously^43^. Mice were euthanized with CO_2_ according to IACUC guidelines, and perfused brains were removed and cut into small pieces before homogenization in 1x Dulbecco’s phosphate-buffered saline with Calcium and Magnesium (DPBS^++^) with 0.4% DNase-I on the tissue dissociator at 37 °C. After filtering the cells through a 70-μm filter, myelin was first removed using Percoll PLUS reagent (GE Healthcare #45001754), then again by using myelin removal beads. After myelin removal, CD11b^+^ cells were selected from the single-cell suspension using the CD11b beads (Miltenyi) as per the manufacturer’s instructions. The CD11b^+^ cells were finally resuspended in microglia growth media^69^, further diluted in TIC (TGF-β, IL-34, and cholesterol containing) media with 2% FBS (Atlanta Biologics #S11150, Lot #H17115) before seeding 1×10^5^ cells per 500 μL in a well of a 24-well plate (Falcon). The cells were maintained in TIC media at 37°C and 10% CO_2_, with media being changed every other day until the day of Aβ treatment (around 12-14 days *in vitro* (d.i.v.)).

### Aβ preparation

The solid human Aβ1-42 peptide (Anaspec #20276) was prepared per our previously-described protocol^70^. Briefly, the peptide was dissolved in 20 mM NaOH, pH=10.5 to make ∼100 μM stock solution. Peptide aggregation was initiated by incubating the solution at 37°C for 24 hours. The peptide was then either stored at −80°C or used directly on the cells after being diluted in culture medium and filtered through a 0.22 μm syringe filter.

### Treatment of Aβ

Primary microglia were treated with 500 μL/well of 500 nM Aβ1-42 or vehicle for 1, 12, or 24 hours. After the respective time points, the conditioned media (CM) from each well was collected and stored at −80°C. The cells were then detached from wells with 0.25% trypsin and collected in 1x phosphate-buffered saline (PBS) before pelleting at 500 x*g* for 6 mins at 4°C. The supernatant was aspirated, and pellets were also stored at −80°C along with the CM for direct injection-MS/MS and MRM-profiling.

### Lipid extraction by the Bligh & Dyer method

Lipid and metabolite extracts were prepared using a slightly modified Bligh & Dyer extraction procedure^71^. The frozen cell pellets from primary microglia were thawed at room temperature, and ultrapure water, methanol, and HPLC-grade chloroform were added to the pellets. The samples were vortexed, resulting in a one-phase solution that was then incubated at 4°C for 15 mins. Next, ultrapure water and chloroform were added, resulting in a biphasic solution. The samples were centrifuged at 16,000 x*g* for 10 mins, resulting in 3 phases in each tube. The bottom organic phase containing the lipids were transferred to new tubes, while the middle phase consisting of proteins and the upper polar phase were discarded. The solvents from the organic phase were evaporated in a speed-vac. The dried lipid extracts were dissolved in acetonitrile/methanol/300 mM ammonium acetate (3:6.65:0.35 v/v/v). The lipid extract solutions were diluted further 50 times before running them on a mass spectrometer.

### Unbiased lipidomic and TAG-species profiling using MRM-profiling

To determine if there were differences in lipid profiles that occur with Aβ activation over 24 hours, lipids extracted from cell lysates as well as conditioned medium were processed for Multiple Reaction Monitoring (MRM)-profiling^72,73^. Instruments used in these experiments are listed in the supplemental material. The detailed methodology of MRM-profiling for targeted lipid profiling has been described previously^74^. This method enabled the interrogation of the relative amounts of numerous lipid species within ten major classes of lipids based upon the LipidMaps database^75^. The lipid classes, along with the total number of MRM transitions screened, are presented in **Fig. S1b**. Due to the large number of triacylglycerol (TAG) lipid species interrogated in this study, TAGs were run in two separate methods (TAG 1 and TAG 2), each measuring relative ion transitions for different TAG species. The TAG species measured in each method were arbitrarily divided. For sample preparation, dried lipid extracts were diluted in methanol:chloroform (3:1 v/v) and injection solvent to obtain a stock solution. Then, the diluted lipid extract was delivered to the ESI source of an Agilent 6410 triple quadrupole mass spectrometer to acquire the mass spectrometry data by flow injection (no chromatographic separation). The raw MS data obtained were processed using an in-house script, and the lists containing MRM transitions along with the respective ion intensity values were exported to Microsoft Excel for statistical analyses to identify the significant lipids and metabolites in Aβ-treated versus vehicle-treated microglia. Individual TAG species were also profiled in primary cultured microglial cells treated with Aβ and vehicle controls, as well as 5xFAD and WT acutely-isolated microglia using MRM-profiling methodology as described above. Briefly, diluted lipid extracts were directly infused into the Agilent Jet Stream ion source of an Agilent 6495C triple quadrupole mass spectrometer. TAG molecular species were identified based upon previously-established MRMs.

### Statistical analyses for lipidomics and metabolomics

Statistical analyses were performed according to our recently-published study^76^. The edgeR package^77^ was used for Aβ-versus vehicle-treated microglia comparisons, as well as comparisons of 5xFAD vs. WT microglia. Abbreviation “ *s*” was used to denote sample, which represent different replicates of an analyte class and “*b*” was used to indicate a biomarker such as a single lipid or metabolite. Ion counts of a biomarker were denoted using these two subscripts. The experimental blank that was done with the injection media was modeled as an ‘intercept’ sample in the analysis to make certain that comparisons were significant with respect to the blank. A generalized linear model was fitted using the edgeR package^77^ for the mean variance as follows:

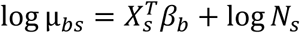

Here, *N*_*s*_ represents the sum of all ion intensities for the sample *s*. The coefficient of variance (CV) for a biomarker ion counts in a sample (*y*_*bs*_) can be calculated using the following equation:

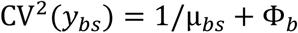

Dispersion of the biomarker was denoted as Φ_*b*_ and it was estimated using the common dispersion method^78^. The associated log-fold change was calculated between the Aβ-treated and vehicle-treated microglia and p-values were obtained using the likelihood ratio test. The BH method was used to calculate p-values to acquire false discovery rates (FDRs)^79^ and a lipid or a metabolite was considered to be significant if fold change > 0.5 and FDR < 0.1.

### Lipid droplet staining of acutely-isolated microglia in suspension

CD11b^+^ cells were isolated from male and female mice (3-4 or 5-7 months old; 5xFAD or WT) and resuspended in 1x PBS, counted using a hemocytometer (1:10 ratio of trypan blue to cell suspension) and stained with 2 μM BODIPY in 1x PBS for 1 hour at 37°C. The cells were then washed once, resuspended in 1x PBS and taken for analysis on an Attune NxT flow cytometer (Invitrogen) after staining of dead cells with DAPI.

### Perfusion and tissue processing

Mice were euthanized with CO_2_ and transcardially perfused with PBS and 4% paraformaldehyde (PFA). Brains were extracted and coronally sectioned (50 µm) using a vibratome and stored in antigel solution (30% glycerol, 30% ethylene glycol, in PBS) at −20°C until use for IHC staining.

### Immunohistochemistry and staining

Free-floating sections were washed five times in PBS, followed by incubation with Methoxy X04 (10µM, Tocris Bioscience,) solution in PBS for 15 minutes. Sections were then stained with HCS LipidTox Green Neutral Lipid Stain (1:1000, ThermoFisher) in PBS for 15 min, followed by incubation with antigen retrieval buffer (10mM sodium citrate, 0.05% Tween20, pH=6.0) at 70°C for 40 min. The sections were allowed to cool, washed, and treated with 0.1% NaBH4 for 30 min. Following antigen retrieval, sections were blocked with blocking buffer (5% NGS, 0.01% Triton X-100, in PBS) for 1 hr at room temperature. The sections were then incubated overnight with the following primary antibodies: anti-IBA1 (1:150, Millipore Sigma) anti-DGAT2 (1:150, ThermoFisher) in blocking buffer at 4°C. Post-incubation, sections were washed thoroughly with PBS + 0.01% Triton X-100 and incubated with secondary antibodies: Goat anti-Mouse Alexa Fluor 594), Goat anti-Mouse Alexa Fluor 647 and Goat anti-Rabbit Alexa Fluor 594 (all from Invitrogen and diluted at 1:500) for 1.5 h at room temperature. Following washes with PBS + 0.01% Triton X-100, sections were mounted on slides, allowed to dry, and coverslipped using Fluoromount-G anti-fade mounting medium (Southern Biotech).

### *Ex-vivo* Lipid droplet staining and Aβ phagocytosis assay of acutely-seeded microglia

Following the protocol described above for cell isolation, 100K CD11b^+^ cells from 5-7-month-old female C57BL/6J mice were seeded in TIC media for 1 hour, followed by 500 nM Aβ^pH^ (in TIC media) / vehicle treatment for 30 minutes. Post-treatment, the cells were stained with LipidTox (1:200) in 1xPBS for 30 minutes at 37°C. After staining, the cells were washed with 1x PBS, detached, and collected in ice-cold 1x PBS. Three minutes before analysis of each sample, DAPI was used to stain dead cells. Single positive controls, gating strategies and analyses were done as described above, but for this experiment live LipidTox^+^ or LipidTox^-^ cells were identified on the Alexa-Fluor 647 channel, while Aβ^pH+^ and Aβ^pH-^ cells were identified on the FITC channel on an Attune NxT flow cytometer (Invitrogen).

### Treatment of DGAT2 inhibitor (D2i) and *ex-vivo* lipid droplet staining and Aβ phagocytosis assay of acutely-seeded microglia

The above protocol was followed with slight modifications for D2i (PZ0233, Sigma) treatment. Briefly, CD11b^+^ cells were seeded TIC media containing 15 µM D2i / vehicle for 1 hour, followed by 500 nM Aβ^pH^ (in TIC media) / vehicle containing 15 µM D2i / vehicle treatment for 30 minutes. Post-treatment, the cells were washed with 1xPBS and stained with LipidTox (1:200) containing 15 µM D2i / vehicle in 1xPBS for 30 minutes at 37°C. After staining, the cells were washed with 1x PBS, detached, and collected in ice-cold 1x PBS containing 15 µM D2i / vehicle and analyzed on the flow cytometer.

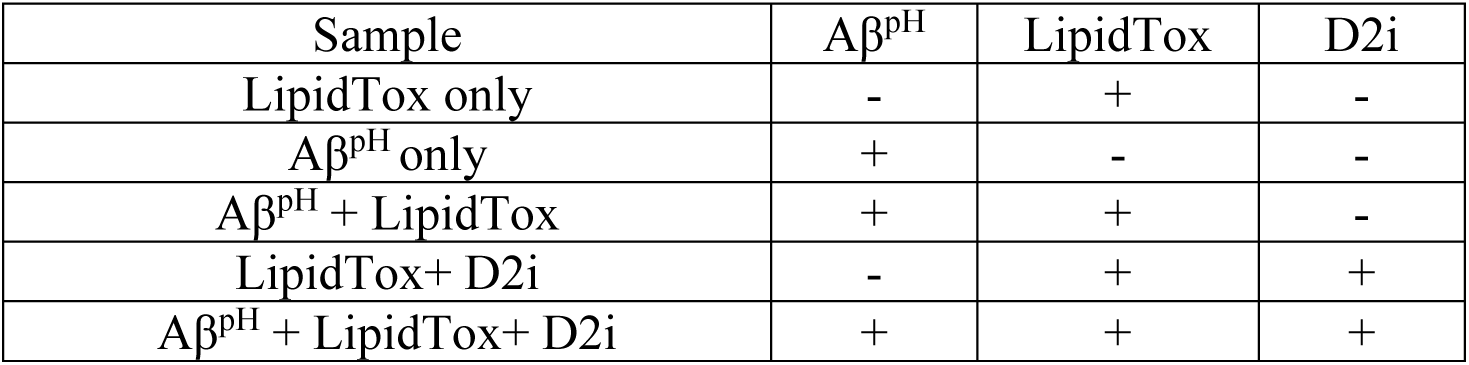

### *Dgat2* mRNA expression by PCR

CD11b^+^ cells isolated from 5-7-month-old female mice (5xFAD and WT) were resuspended in 1x PBS, counted using a hemocytometer (1:10 ratio of trypan blue to cell suspension) and stored at - 80 °C until next step. Total RNA was isolated and purified by using Quick-RNA Miniprep Kit (Zymo Research) following the manufacturer’s protocol. RNA quantification and purity were assessed using Varioskan LUX imaging multi-mode reader (Thermo Scientific). Real-time PCR was conducted using TaqMan probes (Applied Biosystems, Foster City, CA) for Dgat2 (Mm00499536_m1) and the mouse housekeeping gene β-actin (Mm00607939_s1) as an endogenous control. For PCR amplification, an initial denaturation at 95°C for 15 min was followed by 40 cycles of denaturation at 95°C for 10 s and annealing at 60°C for 1 min. Reactions were run in duplicate. Open qPCR software version 1.0.2 (Chai Biotechnologies Inc.) was used for post-amplification analysis. Cq and Tm values were calculated directly by the instrument software and used for finding fold change for *Dgat2* gene expression in 5xFAD relative to WT by the comparative 2^−ΔΔCT^ method.

### Stimulated Raman Scattering (SRS) microscopy for label-free lipid droplet imaging

A dual-output 80-MHz femtosecond pulsed laser source (InSight X3+, Spectra-Physics) was used for the excitation of SRS. The wavelength-tunable output (680-1300 nm) was used as the pump beam and the 1045 nm fixed-wavelength output was used as the Stokes beam. The pump beam was tuned to 800 nm to image CH vibrations in brain samples. Both beams had a pulse duration of ∼120 fs. The Stokes beam was directed into an acousto-optic modulator (ISOMET, M1205-P80L-0.5), which was controlled by a radio frequency driver (ISOMET, 522B-L) and modulated by a function generator (DG1022Z, Rigol). A square wave of 2.5 MHz and 50% duty cycle was used for laser intensity modulation. The 0^th^ order laser beam from the AOM was used for excitation. The beams were combined spatially by a dichroic beam splitter and were chirped using glass rods (SF57, Lattice Electro-Optics). One 150 mm rod was placed only in the probe beam pathway, while two 150 mm rods were used after combining the two beams. We bent the optical beam path to double-pass the two chirping rods to increase the chirping. This gives a 1+4 (Stokes + combined) chirping configuration, which chirps the pump beam to 3.4 ps and the Stokes beam to 1.8 ps. The laser power used on the sample was ∼ 15 mW for the pump and ∼30 mW for the Stokes beam. A motorized linear translational stage (X-LSM050A, Zaber Technology) was used to scan the optical delay between pump and Stokes beams, which were converted to Raman shifts by spectral focusing. The optical delay scanning steps were 10 µm per step. The combined beams were scanned by a 2D galvo scanner set (GVS002, Thorlabs) installed to an upright microscope for imaging. A 60x/1.2 NA water immersion objective lens (UPLSAPO 60X, Olympus) was used to focus the beams onto the sample. The SRS signal was collected by a 1.4 NA oil-immersion condenser. The pump beam was detected with a photodiode detector (S3994, Hamamatsu) with a short-pass filter (980 SP, Chroma technology) to reject the Stokes beam. The alternate voltage signal was amplified using a lab-built tuned amplifier centered at 2.5 MHz. The SRS signal was extracted using a lock-in amplifier (HF2LI, Zurich Instruments). A 2D translation stage (H101, ProScan III, Prior Technology) was used to control sample positions and to perform automated large-area image acquisition and stitching. Data acquisition was enabled using a high-speed data acquisition card (PCIe 6363, National Instruments). The laser scanning and image acquisition was performed by custom-written software based on LabVIEW.

### Analysis of SRS microscopy data

For analysis, the images were saved in .txt files and processed using ImageJ. Pseudocolors were used to display different chemical compositions. Hyperspectral images were used to obtain spectra of lipid droplets and other chemical compositions of the tissue. The spectral profiles were normalized by using a laser intensity profile obtained from cross-phase modulation. For quantitative image analysis a Gaussian blur filter (r=3) was used to process the original image. Then, the processed image was subtracted from the original image to highlight the lipid droplets. Intensity thresholding and particle analysis were then performed using ImageJ built-in functions for quantitative lipid droplet analysis. For quantitative analysis results, areas from the WT and 5xFAD sections were selected and the lipid droplets were analyzed within each area. Merging of different image channels was performed using ImageJ. The percentage was calculated by dividing the number of pixels corresponding to LD signal by the total number of pixels of the entire image.

### Saturated fatty acid structure elucidation using gas-phase ion/ion chemistry

Utilizing gas-phase ion/ion chemistries, the detailed structural elucidation of complex lipids in biological mixtures has been demonstrated previously^80,81^. Here, a charge inversion ion/ion reaction strategy was employed to examine the structure of saturated fatty acids. All experiments were conducted on a Sciex QTRAP 4000 triple quadrupole/linear ion trap mass spectrometer (SCIEX, Concord, ON, Canada) that has been modified to perform ion/ion reactions^82^. To facilitate the mutual storage of oppositely-charged ions, the key instrumental modifications involve the ability to apply AC voltages to the end plates of the q2 reaction cell. Alternately, pulsed nanoelectrospray ionization (nESI) emitters permit the sequential injection of tris-phenanthroline magnesium reagent dications and fatty acid analyte anions^83^. Singly-deprotonated fatty acid anions, denoted [FA – H] anions, generated via direct negative nESI of the lipid extract or authentic reference standard were mass-selected with unit resolution during transient through Q1 and subsequently transferred to the high-pressure collision cell, q2, for storage. Next, positive nESI produced tris-phenanthroline magnesium dications, denoted [MgPhen_3_]^2+^, which were isolated in Q1 prior to accumulation in the reaction cell q2. The [FA – H] anions and [MgPhen3]2+ reagent dications were then mutually stored in q2, yielding the [FA – H + MgPhen_2_]^+^ complex cation. Energetic transfer from the reaction cell q2 to the linear ion trap (LIT), Q3, resulted in the neutral loss of a single phenanthroline ligand and the generation of the charge-inverted complex cation referred to as [FA – H + MgPhen]^+^. Following mass-selection in Q3, the analysis of charge inverted product ions was performed using single-frequency resonance excitation, commonly referred to as ion-trap collision-induced dissociation (CID) (q = 0.383). In summary, reproducible spectral patterns facilitate fatty acid identification^80^. In all cases, mass analysis was performed using mass-selective axial ejection (MSAE)^84^.

### Pathway analysis for metabolomics data

Metabolomic pathway analysis was performed using the MetPA^85^ (metabolomics pathway analysis) tool on MetaboAnalyst 5.0: a free, web-based tool for metabolomics data analysis that uses the KEGG metabolic pathways as the backend knowledge-base. The differentially-regulated metabolites (FDR<0.1) were uploaded into the compound list with hypergeometric test as the enrichment method and relative-betweenness centrality for topology analysis. The KEGG pathway library for *Mus musculus* was chosen as the reference database. All of the matched pathways according to the p values from the pathway enrichment analysis and pathway impact values from the pathway topology analysis were visualized using the “metabolome view” scatter plot.

### Human brain tissue staining

Human hippocampal formalin-fixed paraffin-embedded (FFPE) tissue sections from autopsy samples of both male and female Alzheimer’s disease (AD) patients (>74 years, n=3 per sex) and non-symptomatic (NS) cases (>62 years, n=3 per sex) were used. NS cases were obtained from individuals without any neurological or psychiatric diagnosis, and no chronic systemic inflammatory or infectious condition. All human post-mortem tissue was obtained from the Pathology Research Core in the Robert J. Tomsich Pathology and Laboratory Medicine Institute of the Cleveland Clinic. Institutional ethical guidelines were followed for the appropriate use of these fully de-identified samples for research purposes, after IRB approval. All tissue samples were cut at 15 μm and standard de-paraffinization procedures in xylene and decreasing concentration ethanol solutions were utilized. Sections were stained for amyloid plaques using Amylo-Glo RTD Amyloid Plaque Stain Reagent (Biosensis) according to the manufacturer’s instructions. Then antigen retrieval was performed in 10mM Tris / 1mM EDTA buffer (pH=8.0) for 20 min at 97°C. After cooling down to room temperature, sections were rinsed with distilled water and blocked in 10% normal donkey serum in PBS-Tween-20 0.05% (v/v) for 1 hour. Primary antibodies were added in blocking buffer and sections were incubated for 72hr at 4°C (anti-Adipophilin (PLIN2 Fitzgerald Industries International, 1:200); rabbit anti-DGAT2, (ThermoFisher, 1:200); anti-IBA1, (Millipore, 1:200). Sections were washed with PBS-T 0.05% (v/v) and incubated with secondary antibodies in blocking buffer for 2h at room temperature (Jackson Immunoresearch: donkey anti-guinea pig AF594, 1:500; donkey anti-rabbit FITC, 1:1000; alpaca anti-mouse Cy5, 1:100). Autofluorescence was quenched with TrueBlack-Lipofuscin autofluorescence quencher (Biotium) according to the manufacturer’s instructions and sections were coverslipped with anti-fade fluorescence mounting medium (Abcam). Imaging was performed using a Zeiss LSM 800 confocal microscope using a 40x 1.3NA oil immersion lens and quantification of lipid droplets in relation to microglia and amyloid plaques was performed using the surfaces module in Imaris 9.8.2 (Bitplane).

### Image processing

Confocal microscopy images in Figures **1i**, **2c** and **4b** were processed using Imaris 9.8.2 (Bitplane) to reduce noise by applying the Gaussian or Median filters. For **1i** and **2c**, remaining non-specific speckles in the IBA1 channel (likely produced during the antigen retrieval and immunofluorescence protocol) were removed by size exclusion of 3D objects smaller than 1 or 2 μm^3^ using the Imaris tool "surfaces". A similar approach was also used for confocal microscopy images acquired from human post-mortem FFPE tissue sections that often show artefacts from the deparaffinization and antigen retrieval protocols. 3D objects smaller than 2 or 3 μm^3^ (for IBA1) and 1 μm^3^ (for PLIN2) were removed using Imaris. This size exclusion of speckles and artifacts also ensured that only true LD particles were selected for visualization and quantification in the PLIN2 channel. For the DGAT2 channel, only the Gaussian filter was applied to reduce noise. The mouse brain confocal microscopy images were processed using ImageJ and the despeckle tool was used to remove the fine noise / grainy speckles from the images prior to being used for LD quantification using the ‘Analyze particles’ function coupled with the ROI manager tool.

### *In vivo* administration of DGAT2 degrader and immunohistochemistry

The DGAT2 degrader, synthesized in-house, was dissolved in a mixture of DMSO, propylene glycol, and saline to create a sterile solution. The final solution contained 120 µM of the degrader, 0.6% DMSO, 10% propylene glycol, and 89.4% buffer (0.9% saline solution). Solvents were added to the protein degrader in the precise order specified and thoroughly mixed before proceeding to the next step to avoid any possible precipitation. The DGAT2 degrader or vehicle solutions was then administered to the lateral ventricles of each mouse brain using an intracerebroventricular cannula connected to subcutaneously implanted mini-osmotic pumps (Alzet #2001, 7-day pump, 1µl/hr flowrate). The cannula was implanted at the following coordinates from bregma: −0.5 mm Posterior, −1.1 mm Lateral (Right), and −2.5 mm Ventral (length of catheter)^86^.

After a 7-day treatment with either the DGAT degrader or vehicle, mice were euthanized using CO2 and transcardially perfused with PBS and 4% PFA. Brains were then extracted and coronally sectioned (50 µm thick slices) with a Leica VT1200 vibratome. The sections were stored in an antigel solution (30% glycerol, 30% ethylene glycol in PBS) at −20°C until used for immunohistochemical staining. Coronal brain sections, which included regions of the dorsal hippocampus, were used for analyses. Free-floating sections were washed five times in PBS and then incubated with 2% H_2_O_2_ in 70% methanol for 5 minutes to eliminate autofluorescence. After thoroughly washing them with PBS, the sections were treated with 0.1% NaBH_4_ dissolved in PBS for 30 minutes to remove free aldehyde groups and autofluorescence. After using blocking buffer (10% FBS, 3% BSA, 0.5% Triton X-100 in tris-buffered saline, TBS) for 1 hour at room temperature, the sections were incubated overnight with rabbit anti-IBA1 primary antibody (1:800, Wako # 019-19741) in the blocking buffer at 4°C. Next, the sections were thoroughly washed with TBS + 0.01% Triton X-100 and incubated with goat anti-rabbit Alexa Fluor 488 secondary antibody (Invitrogen, 1:500) for 1.5 hours at room temperature on the following day. After washing with TBS + 0.01% Triton X-100, the sections were incubated with Methoxy X04 (10 µM, Tocris Bioscience) and LD540 (1:500)^87^ in TBS for 15 minutes and mounted on slides, allowed to dry, coverslipped using Fluoromount-G anti-fade mounting medium (Southern Biotech), and imaged using a Zeiss LSM900 confocal microscope. The dorsal hippocampus subicular region was imaged at 20X and used for analyzing LD and Aβ plaques area. The confocal microscopy images were processed using ImageJ for LD quantification using the ‘Analyze particles’ function coupled with the ROI manager tool. The subtract background tool and despeckle tool were utilized to remove the fine noise and grainy speckles from the images before being analyzed.

### Statistical analyses

Data collection was randomized for all experiments and experimenters were blinded for imaging and data analyses. All statistical analyses were performed using GraphPad Prism version 8.2.1 or on R version 4.1.2. Mean between two groups were compared using two-tailed unpaired Student’s t-test. Data from multiple groups were analyzed by one-way analysis of variance (ANOVA) with Tukey’s multiple comparison tests. Information on the sample size, numbers of replicates and statistical test used for each experiment is included in the figure legends.

## Notes

https://github.com/chopralab/microglia_omics

### Summary of Updates

We show that degradation of DGAT2 drastically reduced plaque load and LD abundance in the hippocampus of 5xFAD mice with late-stage AD.

