## Supplementary material for "Amyloid β Induces Lipid Droplet-Mediated Microglial Dysfunction in Alzheimer’s Disease": SI

### SUPPLEMENTARY RESULTS

#### Confirmation of saturated fatty acid structure using gas-phase ion/ion chemistry

Singly-deprotonated fatty acid (FA) anions, generated via direct negative-pulsed nanoelectrospray ionization (nESI), undergo facile charge inversion upon reaction with tris-phenanthroline magnesium $[MgPhen_3]^{2+}$ reagent dications in the gas-phase, yielding abundant $[FA - H + MgPhen_2]^+$ complex cations. Collisional activation of the $[FA - H + MgPhen_2]^+$ complex cation population results in the neutral loss of a single phenanthroline ligand, giving rise to $[FA - H + MgPhen]^+$ as described in the Methods section (**Saturated fatty acid structure elucidation using gas-phase ion/ion chemistry**). The ion-trap collision-induced dissociation CID of the charge-inverted FA complex cation provides rich fragmentation that can be exploited to structurally characterize FAs. For example, ion/ion chemistry has been shown to confirm saturated FA structure in addition to localizing subtle structural features such as the site(s) of unsaturation and cyclopropanation.[85,86,79] Importantly, as ion-trap CID of the $[FA - H + MgPhen]^+$ precursor ion generates CID spectra unique to individual FA isomers, authentic FA reference standards (**Fig. S11a**) can be utilized to confirm FA structure in the case that direct spectral interpretation cannot be used to identify FA structure. In turn, a previously-constructed FA library comprised of ion-trap CID product ion spectra resulting from collisional activation of the $[FA - H + MgPhen]^+$ cations derived from the ion/ion charge inversion of the $[FA - H]^-$ anions of FA standards aids in the identification of FA.

We applied charge inversion ion/ion chemistry to elucidate the structure of FAs 19:0, 20:0, and 22:0 in microglial lipid extracts following Aβ exposure. In brief, collisional activation of saturated FAs of the form $[FA - H + MgPhen]^+$, where FA = 19:0, 20:0, or 22:0, generated a series of product ions spaced evenly at 14 Da apart corresponding to carbon−carbon (C−C) single-bond cleavages along the aliphatic chain (**Fig. S11a**). Even electron, odd mass product ions, began with C3−C4 fragmentation (*m/z* 275) and continued down the aliphatic chain, noting that the only even mass, odd electron product ion (m/z 262) is generated via homolytic C2−C3 cleavage. Interestingly, all CID spectra of charge-inverted saturated FA complex cations described herein display a dominant product ion observed at *m/z* 387. This highly-abundant product ion is most likely obtained via C11-C12 fragmentation in the $[19:0 - H + MgPhen]^+$, $[20:0 - H + MgPhen]^+$, $[22:0 - H + MgPhen]^+$ precursor ions, and is indicative of neutral losses of 114 Da, 128 Da, and 156 Da, respectively (**Fig. S11b**). Explicitly, the individual CID spectra of $[19:0 - H + MgPhen]^+$, $[20:0 - H + MgPhen]^+$, $[22:0 - H + MgPhen]^+$ derived from Aβ-treated microglia (**Fig. S11b**) are in excellent agreement with those obtained from authentic FA reference standards (**Fig. S11a**). Importantly, the utilization of charge-inversion ion/ion chemistry facilitates confident FA identification and confirms structural assignments suggested by MRM experiments.

### SUPPLEMENTARY FIGURES

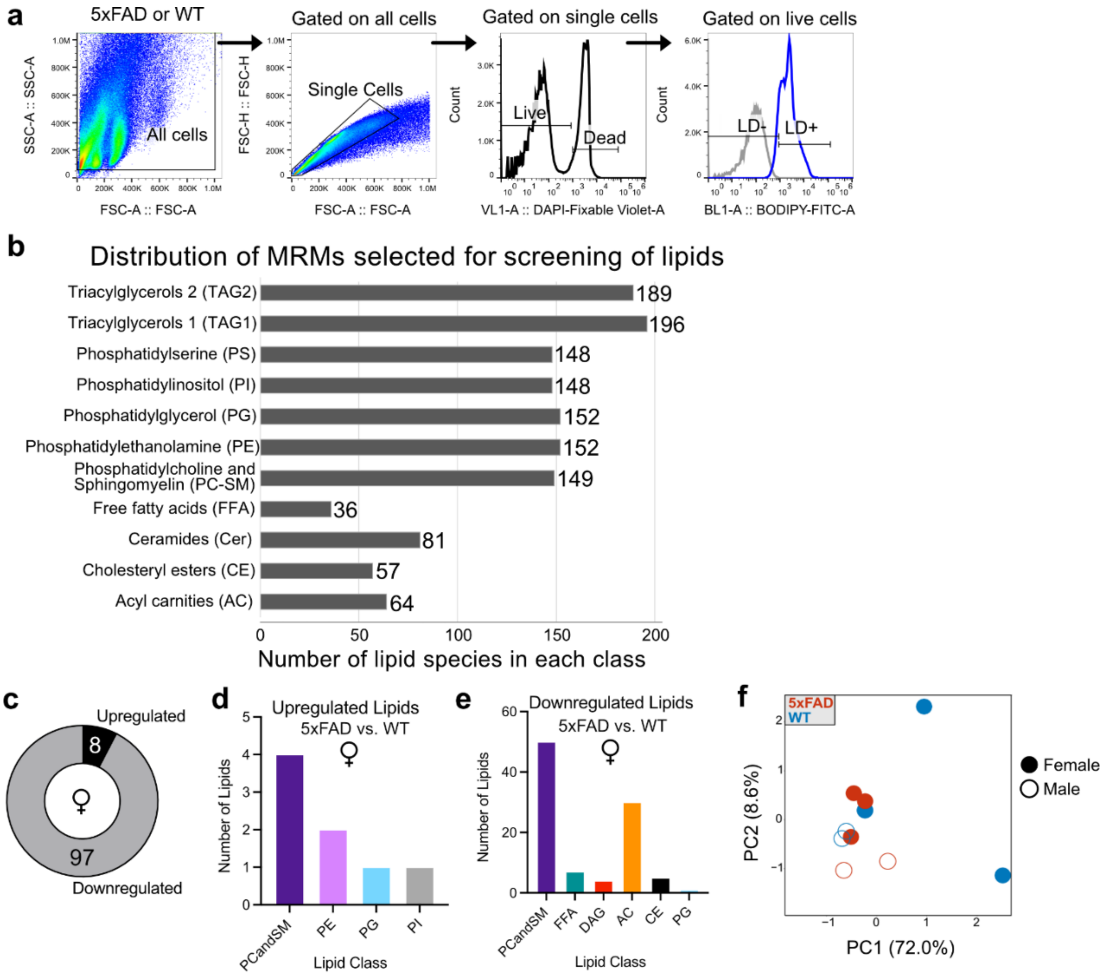


**Fig. S1. LD$^+$ and LD$^-$ microglial subsets from 5xFAD and WT brains can be identified via flow cytometry, and female 5xFAD and WT microglia exhibited unique lipidome characteristics.**

**a.** Gating strategy of flow cytometry analysis to determine LD$^+$ (BODIPY$^+$) and LD$^-$ (BODIPY$^-$) cell subsets among microglia isolated from mouse brains; a representative example from a 5-7-month-old female WT mouse is shown. **b.** Distribution of MRM transitions selected for screening lipids. A total of over 1370 transitions (used to ID lipid species) were organized into 10 MRM-based mass spectrometry methods for lipid classes (TAG1 and TAG2 were combined as the TAGs class). **c.** Direct comparison of lipidomic profiles revealed 8 upregulated and 97 downregulated lipids in female 5xFAD compared to WT microglia. **d.** Upregulated lipid classes (in 5xFAD vs. WT) identified in female microglia: phosphatidylcholine and sphingomyelin (PC-SM) were the lipid class with the highest number of differentially upregulated lipid species, followed by phosphatidylethanolamine (PE), phosphatidylglycerol (PG), and phosphatidylinositol (PI). **e.** Downregulated lipid classes (in 5xFAD vs. WT) identified in female microglia: PC-SM and acyl carnities (AC) were the lipid classes with the highest number of differentially downregulated lipid species. FFAs were also downregulated. **f.** PCA plot showing the distribution of male and female

5xFAD and WT lipidomes. The plot shows greater separation (indicative of their variation) between the female 5xFAD and WT lipidomes (filled circles) compared to the corresponding male lipidomes (unfilled circles) that are closer together, implying less overall variation among them.

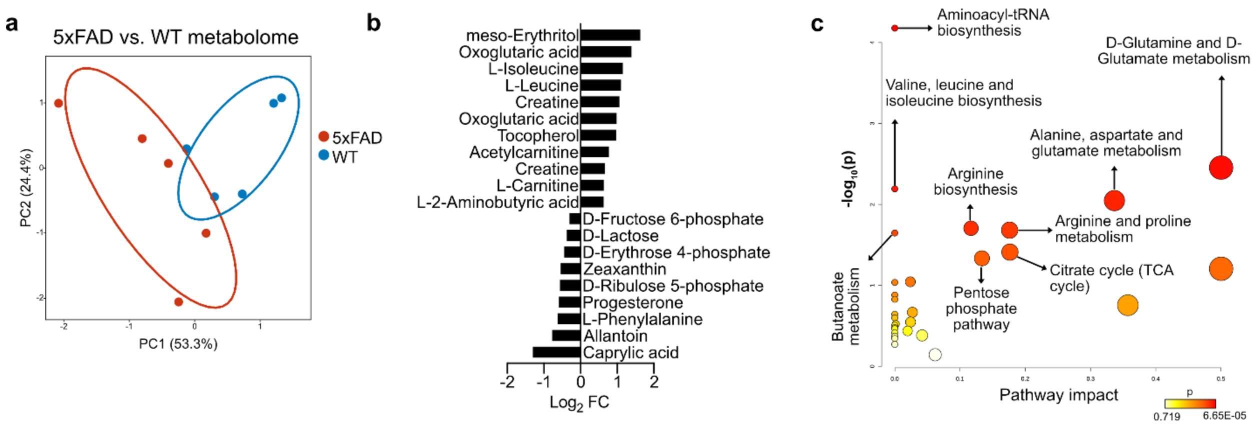


**Fig S2. MRM-profiling reveals changes to metabolite profiles of 5xFAD and WT microglia.** **a.** Direct comparison of the metabolomic profiles between 5xFAD and WT microglia reveals a clear separation in the PCA space, indicative of extensive variation between them. **b.** Plot demonstrating the significantly (FDR<0.1) upregulated ($log_2FC>0$) and downregulated ($log_2FC<0$) metabolites identified in 5xFAD vs. WT microglia. **c.** Pathway analysis of the significantly altered metabolites in 5xFAD vs. WT microglia revealed several interesting cellular pathways that are affected in the chronic AD mouse model, including citrate (TCA) cycle, arginine synthesis, and glutamate metabolism, which is an additional indication of the profound overall metabolic differences among 5xFAD and WT microglia.

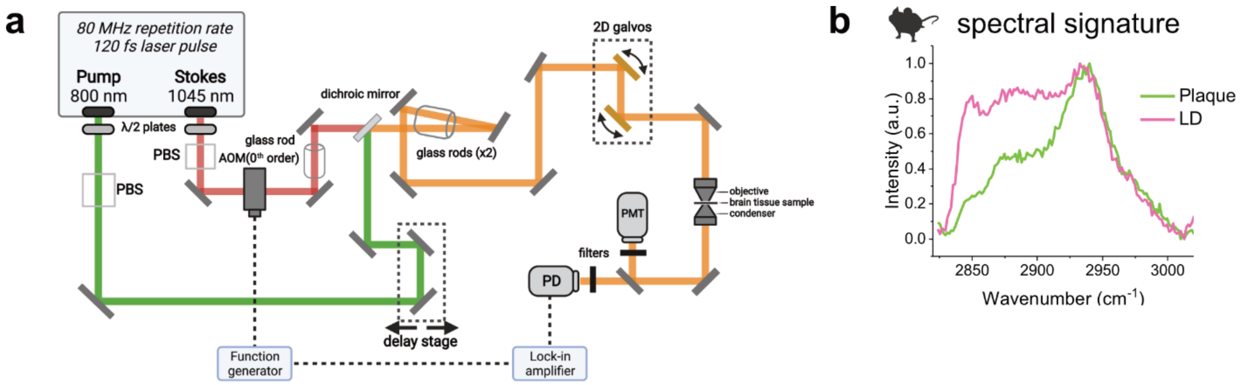


**Fig. S3. Amyloid plaques and lipid droplets are identified with their spectral signature using hyperspectral stimulated Raman scattering (SRS).**

**a.** Schematic of the multimodal microscopy system. A fixed output at 1045 nm and a tunable output set at 800 nm were used for signal generation. Imaging beams were spectrally chirped using glass rods for hyperspectral Raman imaging. SRS signals were collected using a photodiode and demodulated using digital lock-in amplification. A photomultiplier tube was used to collect two-photon excitation fluorescence (TPEF) signals. (λ/2 plate: half-wave plate; PBS: polarization beam splitter; PD: photodiode; PMT: photomultiplier tube; AOM: acousto-optic modulator). **b.** Representative spectral signatures for lipid droplets (magenta, identified using CH2 stretch at 2855 $cm^{-1}$) and plaques (green, correlated with CH3 stretch at 2930 $cm^{-1}$) as they were detected in hippocampal brain slices from 5xFAD mice, using the imaging setup shown in **a**.

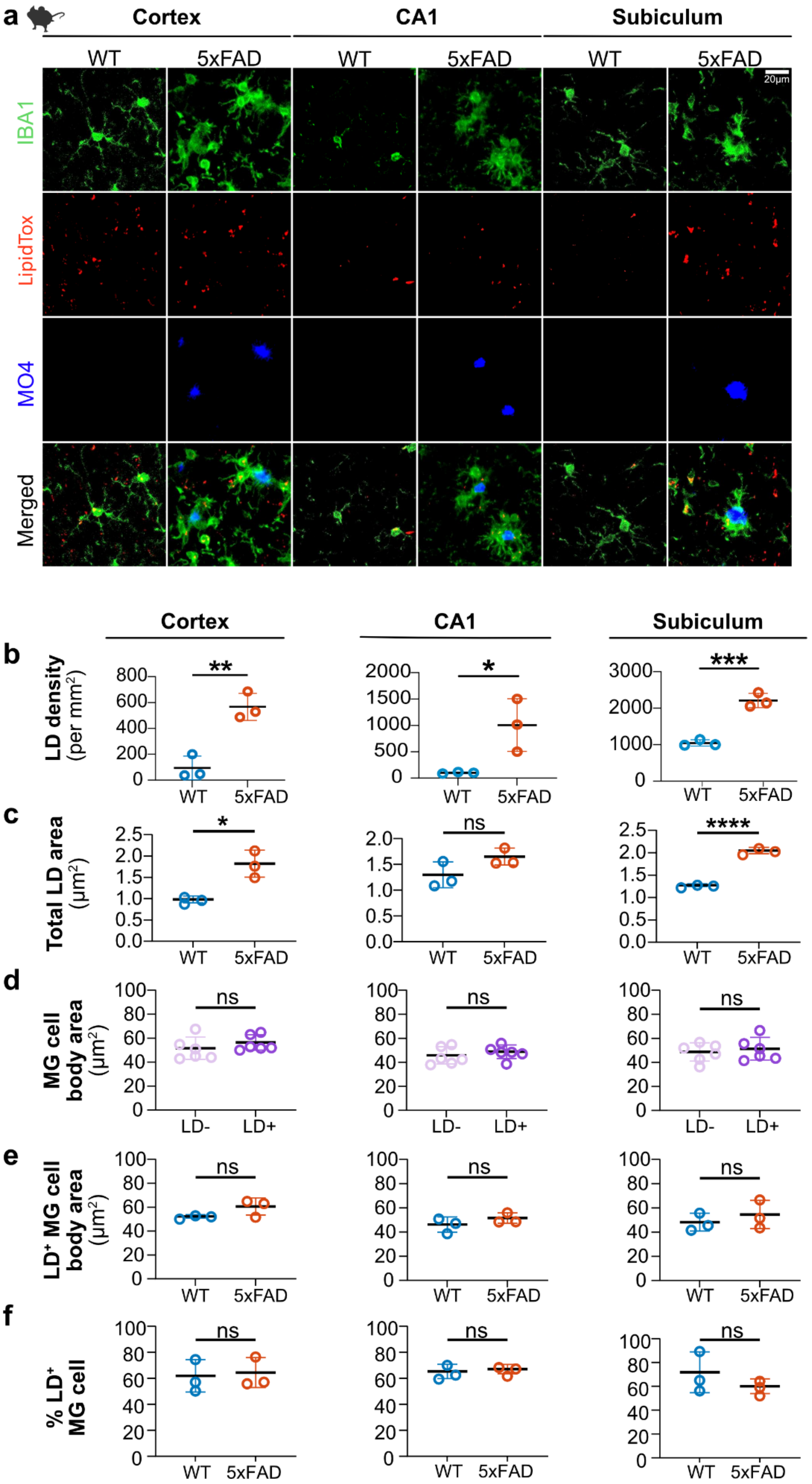
a
Cortex
CA1
Subiculum
WT
5xFAD
WT
5xFAD
WT
5xFAD
20μm
IBA1
LipidTox
MO4
Merged
Cortex
CA1
Subiculum
b
LD density (per mm²)
**
*
***
WT
5xFAD
c
Total LD area (μm²)
*
ns
****
WT
5xFAD
d
MG cell body area (μm²)
ns
LD-
LD+
e
LD⁺ MG cell body area (μm²)
ns
WT
5xFAD
f
% LD⁺ MG cell
ns
WT
5xFAD

**Fig. S4. Lipid-laden microglia in the 5xFAD brain show regional differences in LD density and area.**

**a.** Staining of IBA1 (microglia), LipidTox (LDs), and Methoxy XO4 (MO4; Aβ plaques) in 5xFAD and WT female mouse brain tissue. Representative images from cortex, CA1, and subiculum regions are shown. **b.** Quantification of LD density (per $mm^2$) revealed significantly higher LD density in microglia in cortex, CA1, and subiculum from 5xFAD brains compared to WT; ***P*= 0.0042*,* **P*= 0.0349*,* ****P*= 0.0007. **c.** Quantification showed significantly higher total LD area ($\mu m^2$) in cortex and subiculum from 5xFAD compared to WT mice; **P*= 0.0113*,* *****P*= 0.00006. **d.** Quantification of microglial cell body area ($\mu m^2$) in cortex, CA1, and subiculum from 5xFAD and WT mice showed no significant differences. **e.** Quantification of $LD^+$ microglial cell body area ($\mu m^2$) in cortex, CA1, and subiculum from 5xFAD and WT mice showed no significant differences. **f.** Quantification of % $LD^+$ microglial cells in cortex, CA1, and subiculum from 5xFAD and WT mice showed no significant differences. For **b-f**, data represent mean ± SD. Unpaired t-tests, N=3 mice per group (WT and 5xFAD).

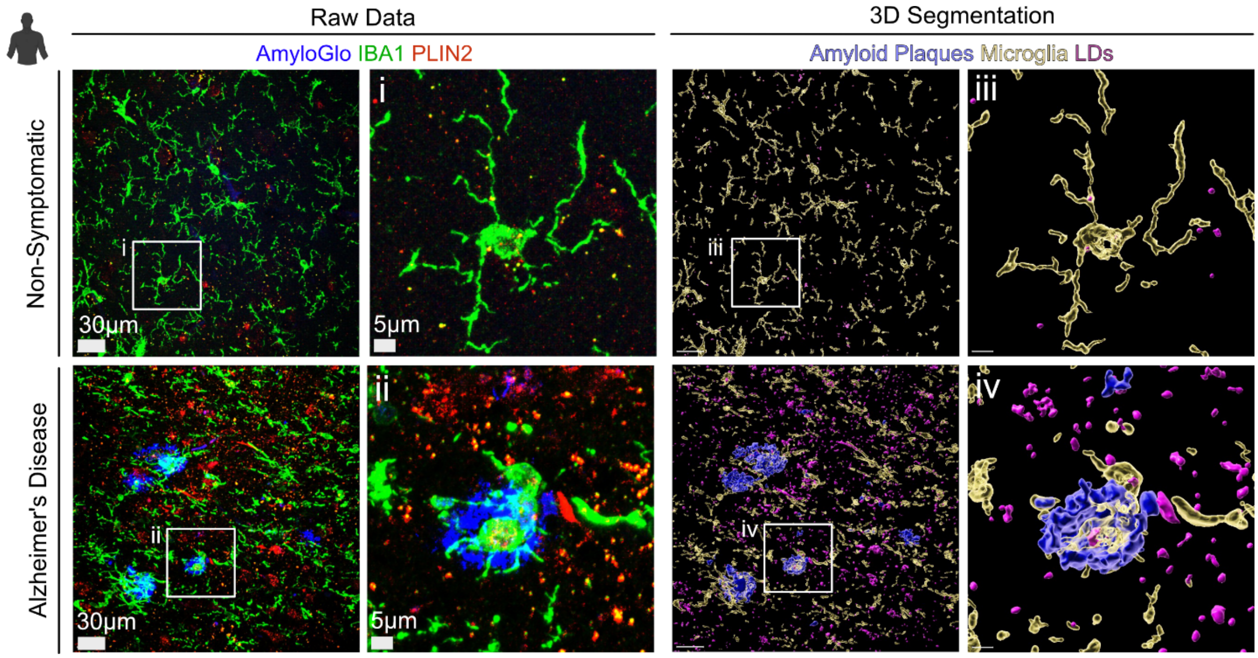


**Fig. S5. 3D reconstruction and segmentation approach for the quantification of LDs in relation to microglia and amyloid plaques on human hippocampal tissue.**

3D reconstruction and segmentation of IBA1, PLIN2, and AmyloGlo immunofluorescent signal on human hippocampal tissue sections from AD and non-symptomatic cases using Imaris (Oxford Instruments). Human FFPE tissue sections (15µm thick) were immunostained for IBA1 (microglia), PLIN2 (LDs), and AmyloGlo (Aβ plaque). Projections from raw confocal microscopy z-stacks of images are shown on the left (entire fields of view and boxed areas) and the equivalent 3D segmentation images produced by using the "Surfaces" tool of Imaris are shown on the right. Scale bars in segmented images are the same as the raw data images (left: 30µm, right: 5µm).

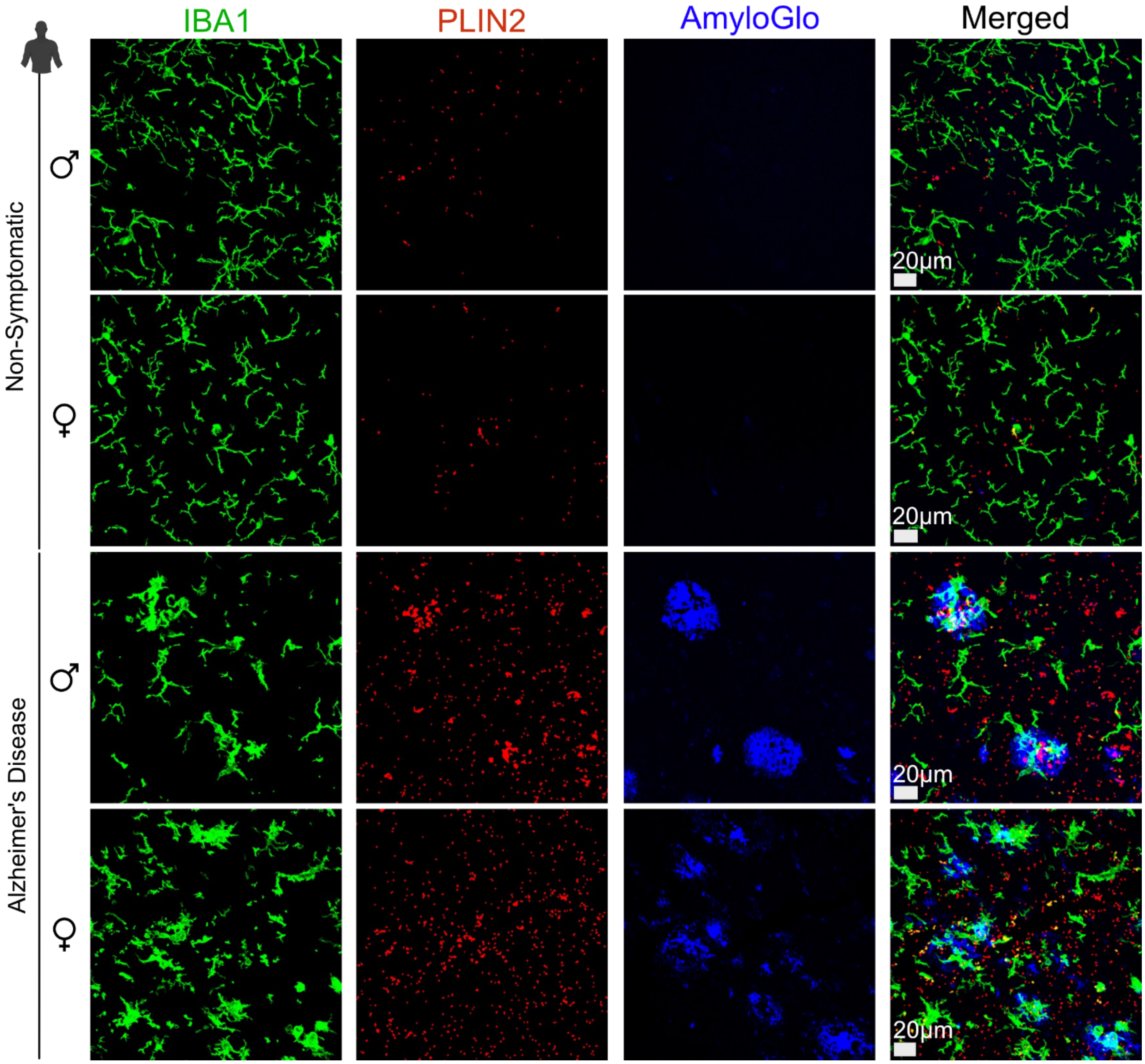


**Fig. S6. Increased LDs and LD⁺ microglia were detected in post-mortem human hippocampal tissue from both male and female AD patients.**

Representative images showing immunolabeled microglia (IBA1), LDs (PLIN2), and amyloid plaques (AmyloGlo) on human hippocampal tissue from male and female AD and non-symptomatic patients. Individual and merged channels are shown for IBA1, PLIN2, and AmyloGlo. Increases in LDs and PLIN2$^+$ LDs in reactive microglia surrounding Aβ plaques in both male and female AD patients were evident.

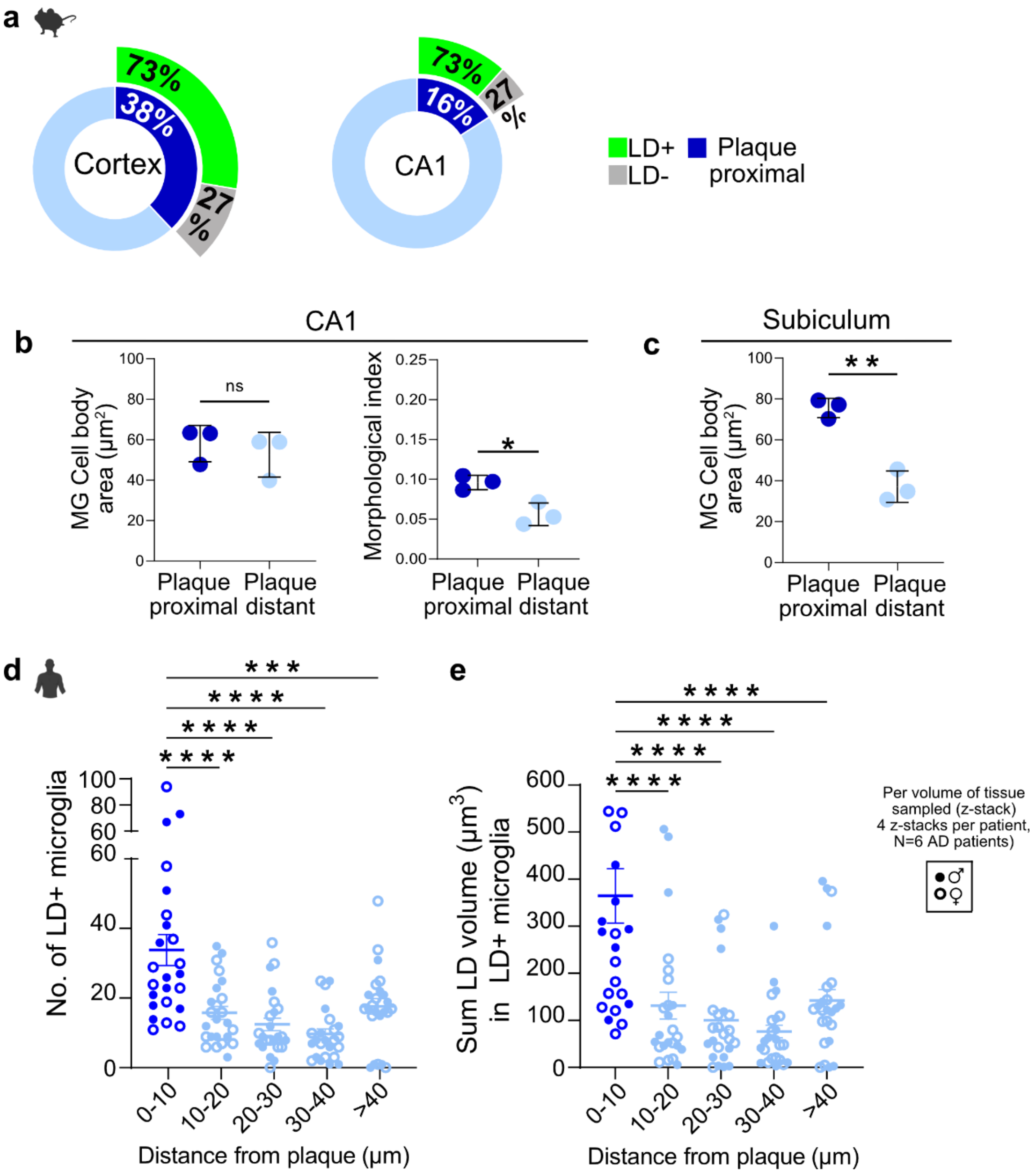


**Fig. S7. In-depth quantification of LDs in mouse and human brain tissue reveals region-specific and plaque-associated increases in microglial LD load.**

**a.** Quantification of % $LD^+$ microglia that were plaque-proximal or -distant in the cortex and CA1 of female 5-7 months old 5xFAD mice. In the cortex, out of all microglia, 38% were in contact with plaques, while 62% were away from plaques. Out of all plaque-contacting microglia, 73% were $LD^+$, whereas only 27% of the plaque-contacting microglia were $LD^-$. Similarly, in the CA1 region, 73% of the plaque-contacting microglia (16% of total microglia) were $LD^+$. **b.** Quantification of microglial cell body area ($\mu m^2$) and microglial morphological index (Cell soma area/Total cell area)[76] between plaque-distant and plaque-proximal microglia in the CA1 region of the hippocampus from the same animals; $*P= 0.0145$. **c.** Quantification of microglial cell body area between plaque-distant and plaque-proximal microglia in the subiculum region of the hippocampus from the same animals. Significantly higher cell body area was observed in plaque-proximal microglia compared to plaque-distant microglia, indicating a unique cellular morphology affected by the Aβ plaque in this region; $**P= 0.0018$. For **b** and **c**: Data represent mean ± SD. Unpaired t-test, N=3 mice per group (WT and 5xFAD). **d.** Quantification of the number of $LD^+$ microglial fragments as a function of distance from the closest amyloid plaque in the hippocampus from human AD patients. Significantly higher numbers of $LD^+$ microglial fragments were observed closer to the plaques (0-10 µm) compared to 10-20µm (****$P$= 0.00002735), 20-30µm (****$P$= 0.00000049), 30-40µm (****$P$= 0.00000001) and >40µm (***$P$= 0.0002085) from the plaques. **e.** Quantification of the sum of LD volume in $LD^+$ microglial fragments as a function of distance from the closest plaque in the same human AD brain tissue. A significantly higher LD volume was observed closer to the plaques (0-10 µm) compared to 10-20µm (****$P$= 0.00001323), 20-30µm (****$P$= 0.00000067), 30-40µm (****$P$= 0.00000006) and >40µm (****$P$= 0.00003707) from the plaques. For **d** and **e**: Data represent mean ± SEM. One-way ANOVA with Tukey's multiple comparison tests, values represent each confocal z-stack of the tissue imaged (4 z-stacks per patient), from a total of 6 AD patients (3 male and 3 female).

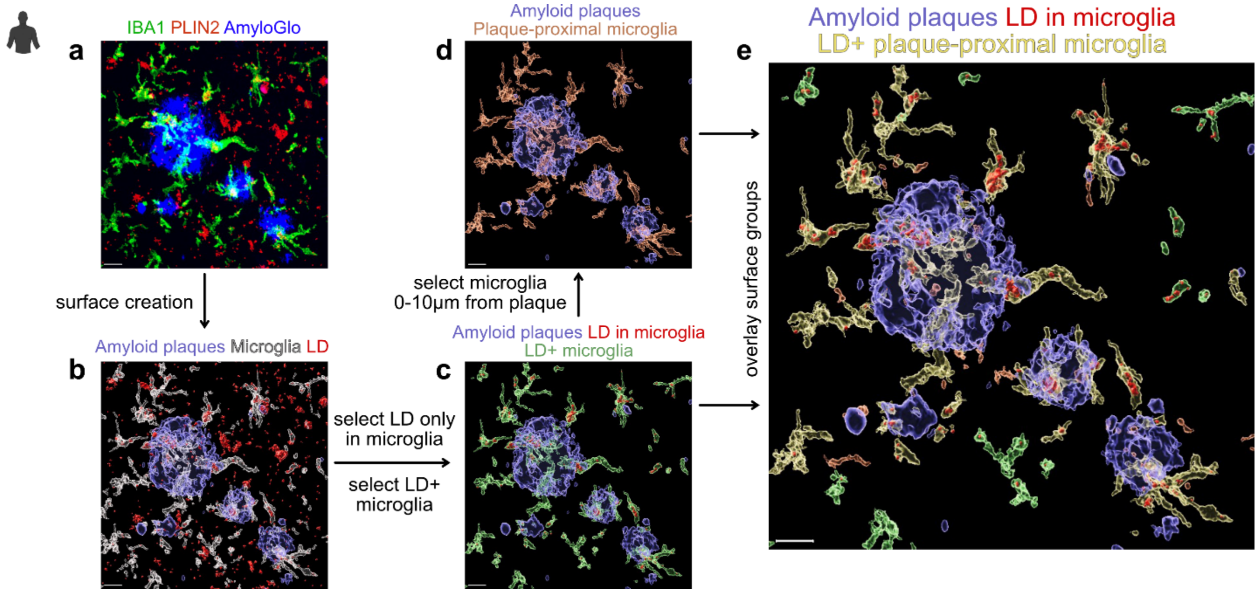


**Fig. S8. Step-by-step workflow for the quantification of LDs in 3D in plaque-proximal and plaque-distant microglia from immunostained human hippocampal tissue.**
Schematic description of 3D volumetric analyses of LDs in plaque-proximal and plaque-distant microglia from human hippocampal FFPE tissue stained for IBA1/PLIN2/AmyloGlo using Imaris. The first step was the creation of 3D "surfaces" for each channel of the 3D reconstructed confocal z-stacks (**a, b**). Then, for the analysis of LD volume within $LD^+$ microglia, only the microglial "surfaces" containing LDs ($LD^+$ microglia) were selected (**c**). Then, among those $LD^+$ microglia, only the microglia that were proximal to the plaques (0-10µm distance from closest plaque) were selected (**d**). Finally, the quantification was performed for both plaque-proximal microglia (0-10µm from closest plaque) and plaque-distant microglia (10-20, 20-30, 30-40, >40 µm from closest plaque) (**e**) and was reported in the corresponding figures as a function of that distance. Scale bar: 15µm.

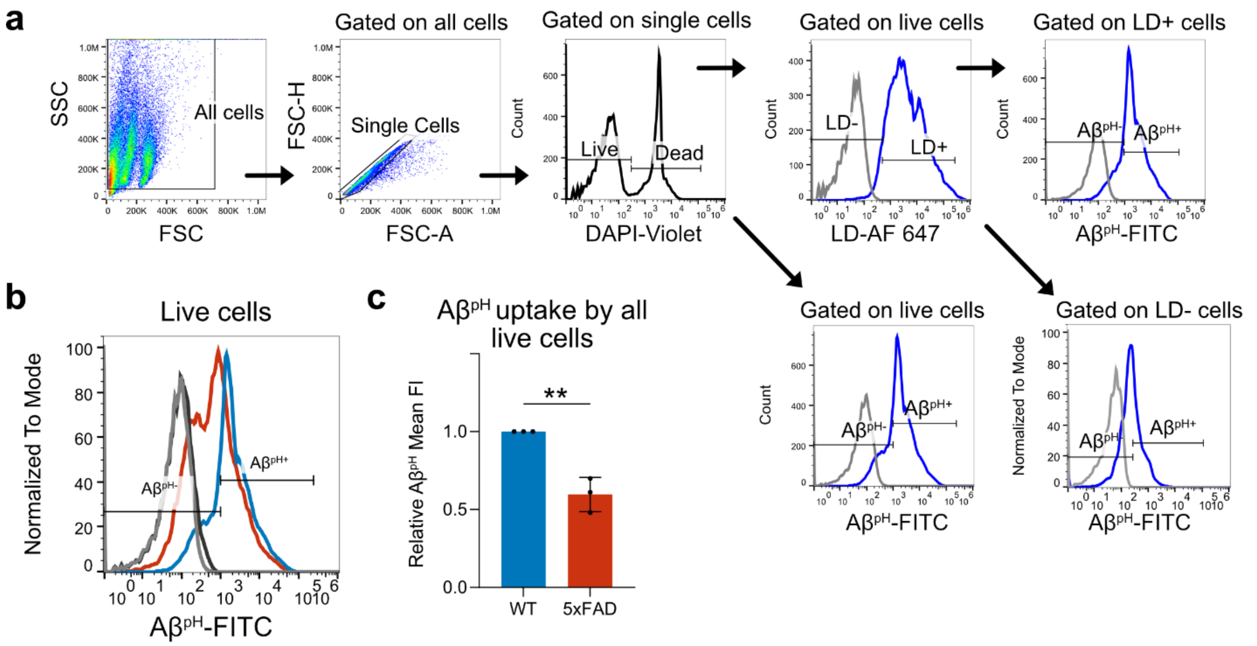


**Fig. S9. Gating strategy and quantification of $A\beta^{pH}$ uptake as a measure of the phagocytic capacity of acutely seeded 5xFAD and WT microglia.**

**a.** Representative gating strategy for the flow cytometry analysis of $A\beta^{pH}$ phagocytosis by $LD^{+}$, LD-, as well as all live ($DAPI^{-}$) microglia; data shown here is from acutely isolated $CD11b^{+}$ cells pooled from whole brains of three 5-7-month-old female WT mice. **b.** Histogram showing $A\beta^{pH+}$ and $A\beta^{pH-}$ microglial cell subpopulations from all live cells in 5xFAD (red) and WT (blue) brains. 5xFAD microglia showed reduced $A\beta^{pH}$ uptake compared to WT microglia. Unstained controls for WT and 5xFAD shown as light grey and dark grey (solid) lines, respectively. **c.** Quantification of $A\beta^{pH}$ uptake by 5xFAD relative to that by WT microglia from the entire live population. 5xFAD microglia exhibited significantly reduced $A\beta^{pH}$ phagocytic capacity compared to WT microglia; *$**P=0.0032$.* Data represent mean ± SD. Unpaired t-test, cells were pooled from 3 mice per group (3 WT and 3 5xFAD mice) for each of the N=3 experiments.

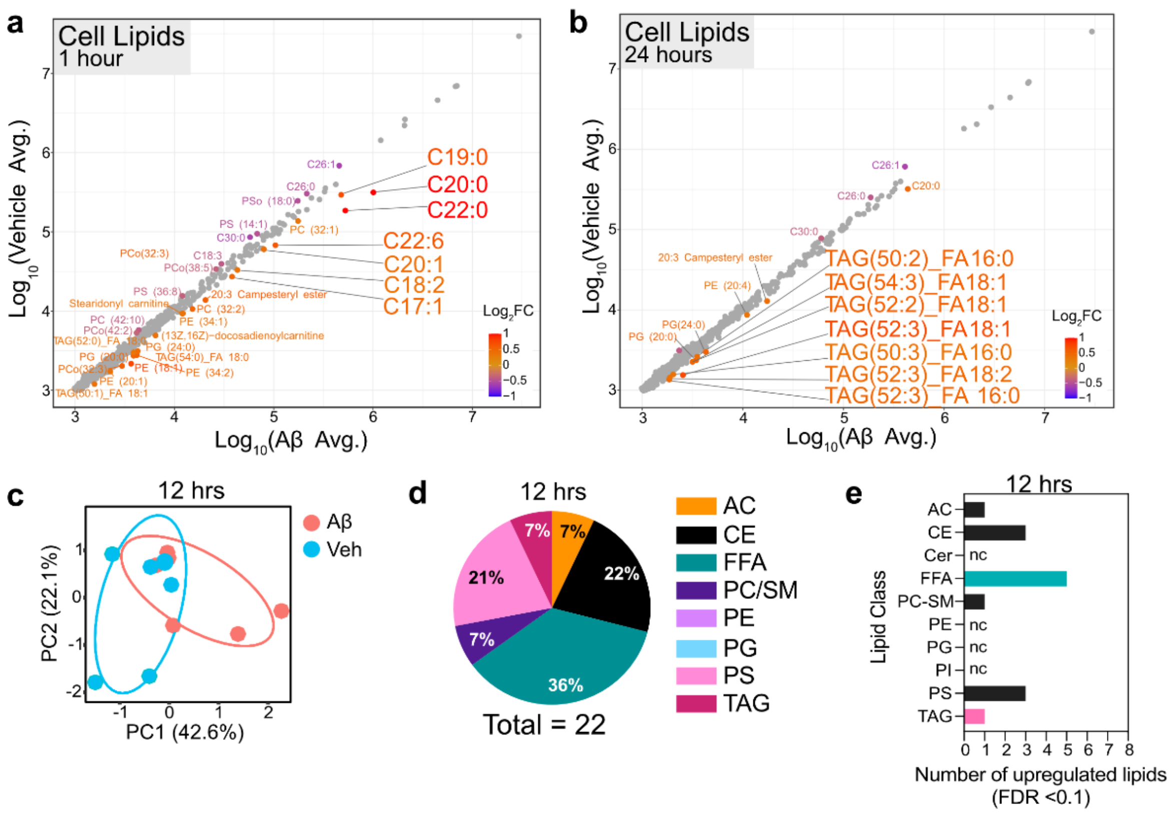


**Fig. S10. Multiple Reaction Monitoring-profiling of 1349 lipid species and 717 metabolites reveals profound changes to microglial lipid and metabolite profiles upon Aβ treatment.**
**a.** Scatter plot showing the differentially-regulated lipids between Aβ and vehicle-treated microglia at the 1-hour time point. Select long-chain saturated FFAs (especially C19:0, C20:0, and C22:0) upregulated in Aβ-treated microglia are highlighted. **b.** Scatter plot showing the differentially-regulated lipids between Aβ vs. vehicle-treated microglia (cells) at the 24-hour time point. The individual TAG species upregulated in Aβ-treated microglia are highlighted. **c.** PCA plot that shows the extensive variation between the overall lipidomic profiles of microglia after 12 hours of treatment with Aβ compared to vehicle-treated cells. **d.** The distribution of significantly-regulated lipid classes identified in microglia at 12 hours of Aβ treatment. 22% and 36% of the differentially-regulated lipids at 12 hours were CEs and TAGs, respectively – both neutral lipids found within LDs. These data indicate a pathway towards an LD-rich metabolic shift in microglia between 1 and 24 hours of Aβ treatment. **e.** Upregulated lipid classes at 12 hours of Aβ treatment. Lipidomics data are from N=5, 7-month-old male and female mouse brains i.e., cells from each brain were cultured independently and treated with vehicle or Aβ. Cell pellets and conditioned media were collected for the lipidomics experiment.

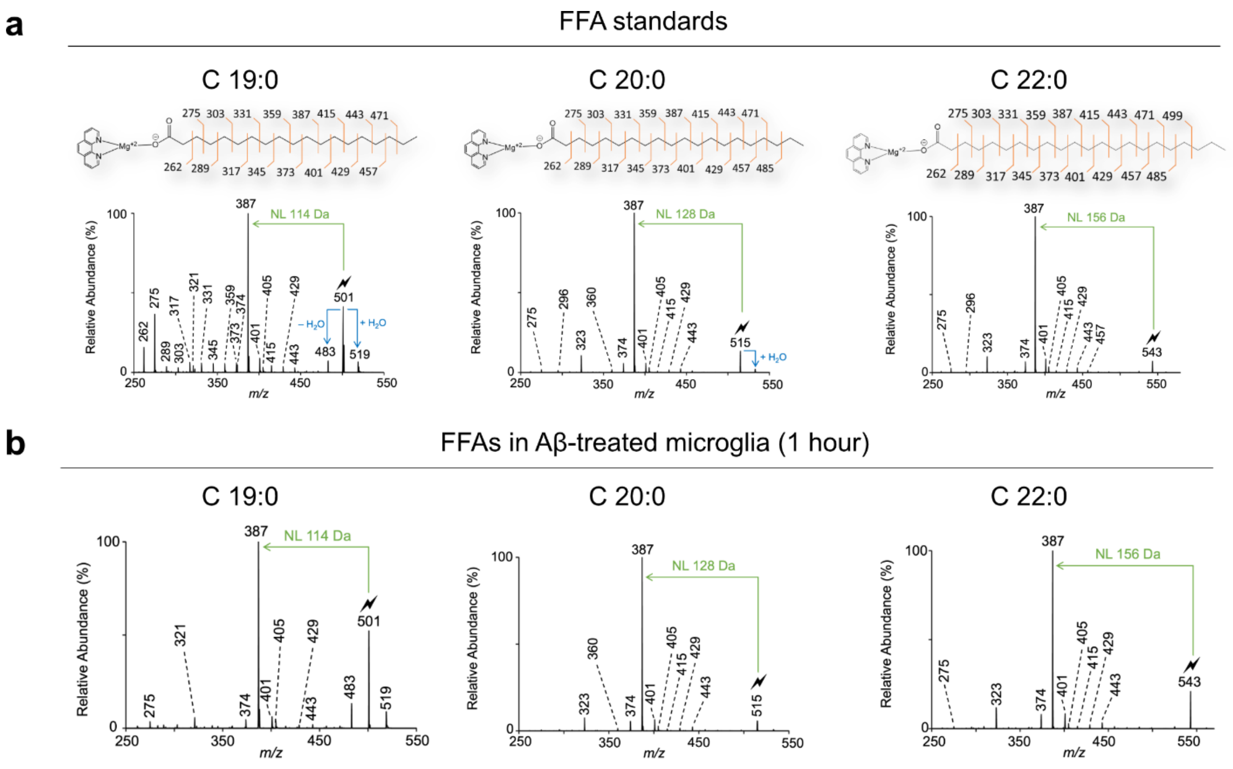


**Fig. S11. Gas-phase ion/ion chemistry validated the chemical structures of the top differentially altered long-chain saturated FFAs that were identified by MRM-profiling.**
**a.** Spectral plots corresponding to the lipid standards used for validation of cellular lipids. Neutral losses (NL) of 114, 128, and 156 Da were observed in C19:0, C20:0, and C22:0, respectively. **b.** Structural verification of FFA species C19:0, C20:0, and C22:0 in the Aβ-treated microglia. The NL observed in each FFA spectrum corresponds to the NL observed in the standard plots, thereby verifying the structural identity of the individual FFA species in the examined microglial samples.

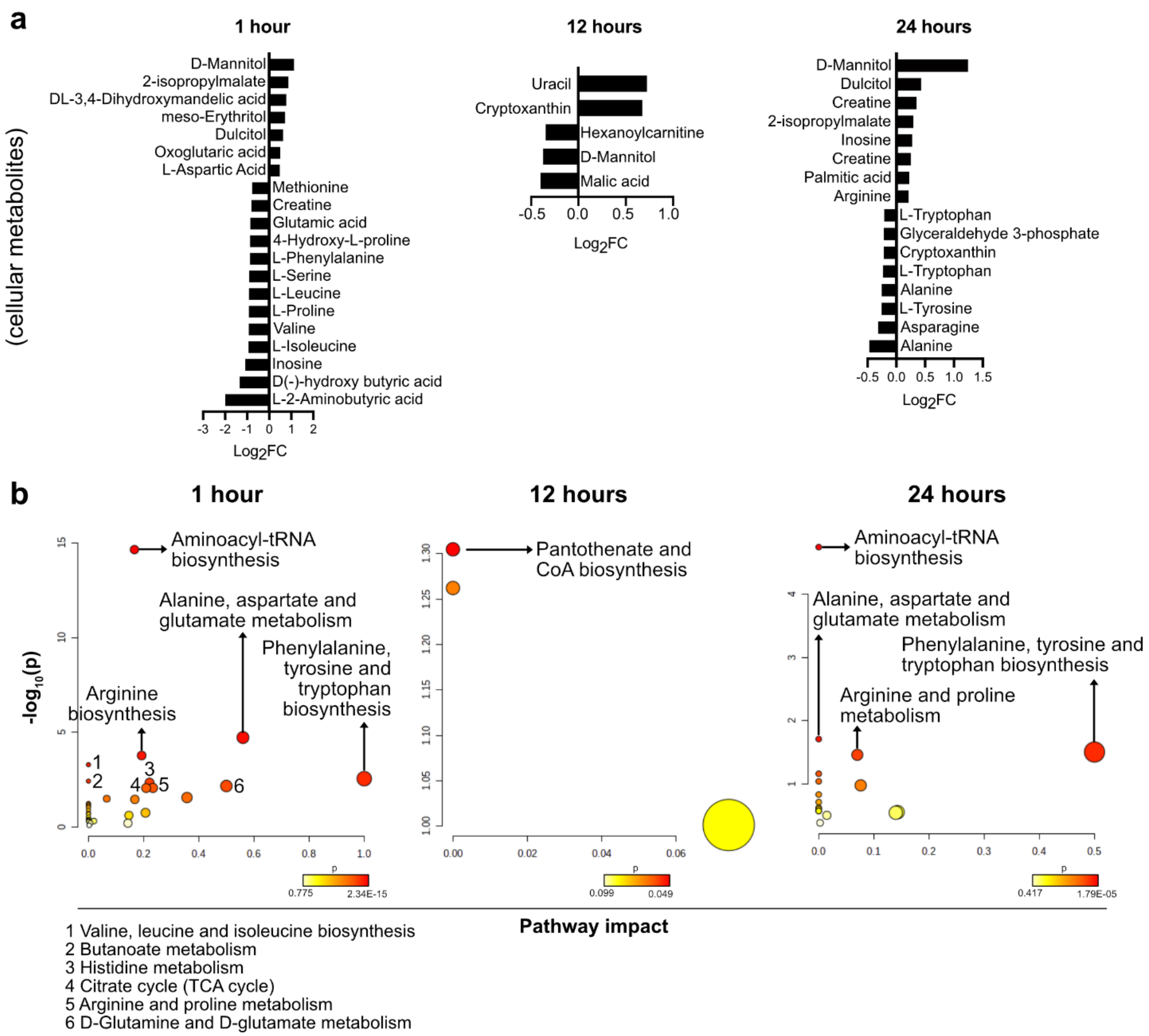


**Fig. S12. Aβ causes significant changes to microglial intracellular metabolites and related pathways after 1, 12, and 24 hours of treatment.**

**a.** Profiling of intracellular metabolites from microglia after 1, 12, and 24 hours of Aβ treatment. Plots demonstrate the significantly (FDR<0.1) upregulated ($log_2FC>0$) and downregulated ($log_2FC<0$) metabolites in microglia at each time point of Aβ treatment, compared to vehicle treated cells. **b.** Pathway analysis of the significantly altered metabolites at 1, 12, and 24 hours of Aβ treatment revealed several interesting cellular pathways that were affected due to Aβ exposure, such as arginine metabolism and alanine/aspartate/glutamate metabolism.

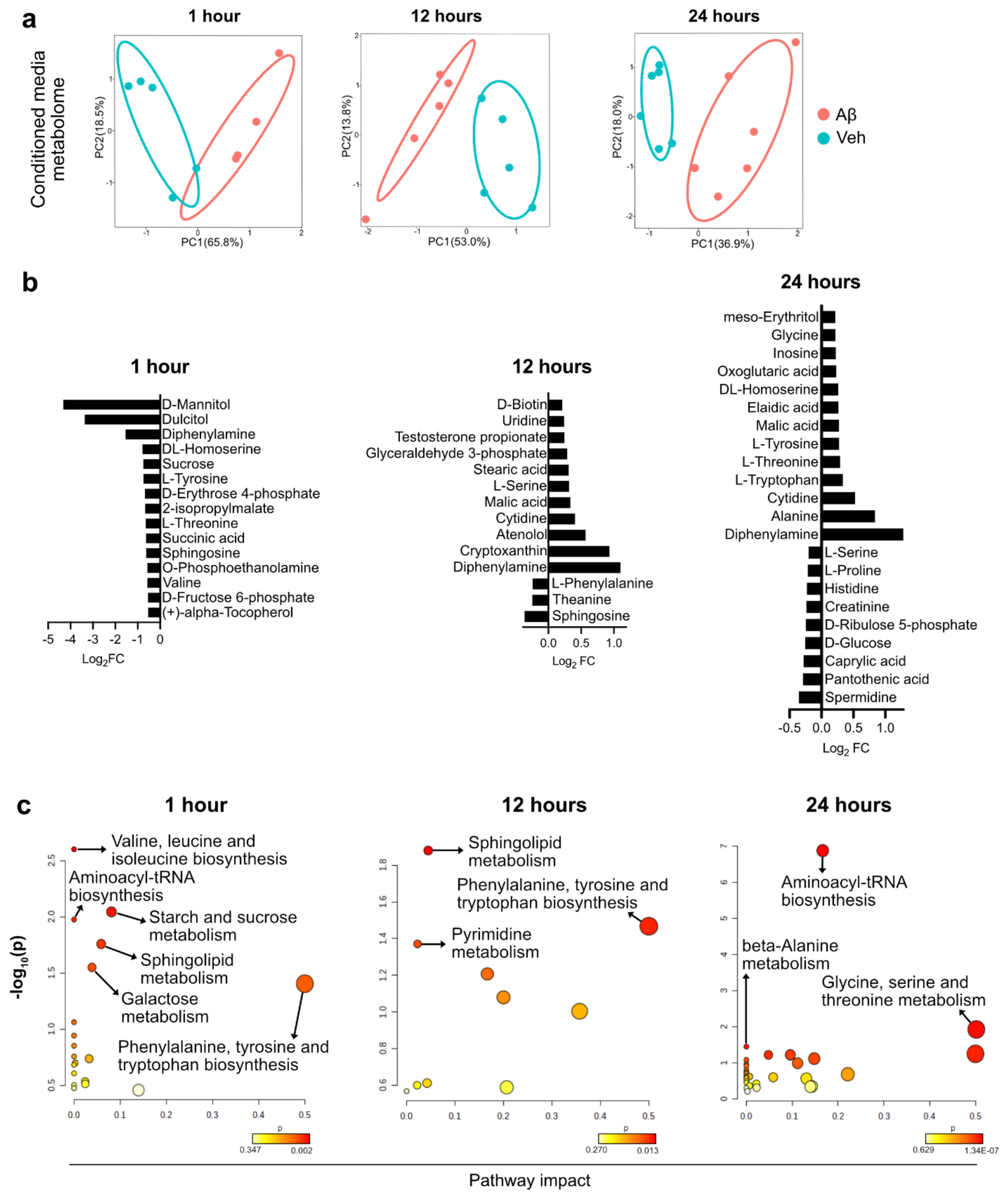


**Fig. S13. Aβ causes significant changes to microglial secreted metabolites and related pathways after 1, 12, and 24 hours of treatment.**

**a.** Profiling of secreted metabolites from microglia (collected from conditioned media) after 1, 12, and 24 hours of Aβ treatment. PCA plots show clear separation (significant variation) between the microglial metabolomes of cells treated with Aβ for1, 12, and 24 hours compared to vehicle-treated cells. Overall, these plots demonstrate the variance observed in 84.3%, 66.8%, and 54.9% of the data at 1, 12, and 24 hours, respectively. **b.** Plots demonstrating the significant ($FDR<0.1$)

upregulated ($log_2FC>0$) and downregulated ($log_2FC<0$) metabolites identified in microglial conditioned medium at each time point of Aβ treatment. There were no upregulated metabolites after 1-hour of Aβ treatment, compared to vehicle. **c.** Pathway analyses of the significantly altered metabolites in the microglial conditioned medium at 1, 12, and 24 hours of Aβ treatment revealed several cellular pathways that were affected due to increasing Aβ exposure, compared to vehicle-treated cells.

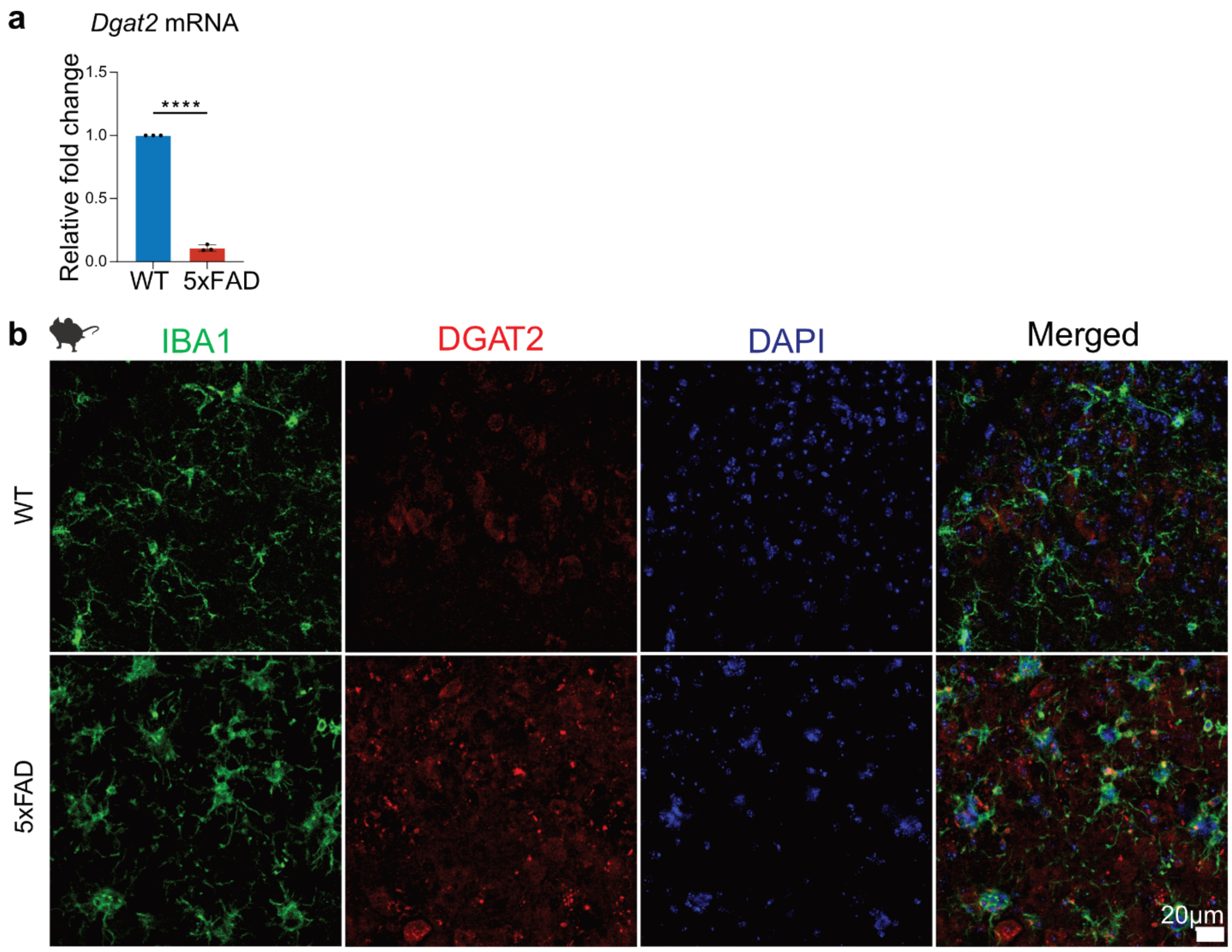


**Fig. S14. *Dgat2* mRNA in microglial cells is decreased but the DGAT2 protein is higher in 5xFAD compared to WT brains.**

**a.** Quantification of microglial *Dgat2* mRNA expression by qPCR. *Dgat2* mRNA was decreased in 5xFAD compared to WT microglia; ****$P<0.0001$. Data represent mean ± SD. Unpaired t-test, cells were pooled from 3 mice per group (3 WT and 3 5xFAD mice) for each of the N=3 experiments. **b.** Staining of IBA1 (microglia), DGAT2, and DAPI (nuclei) in 5xFAD and WT mouse brain tissue. Representative images of the subiculum region are shown. N=3 female mice per group (WT or 5xFAD). Quantification is shown in figure 4b.

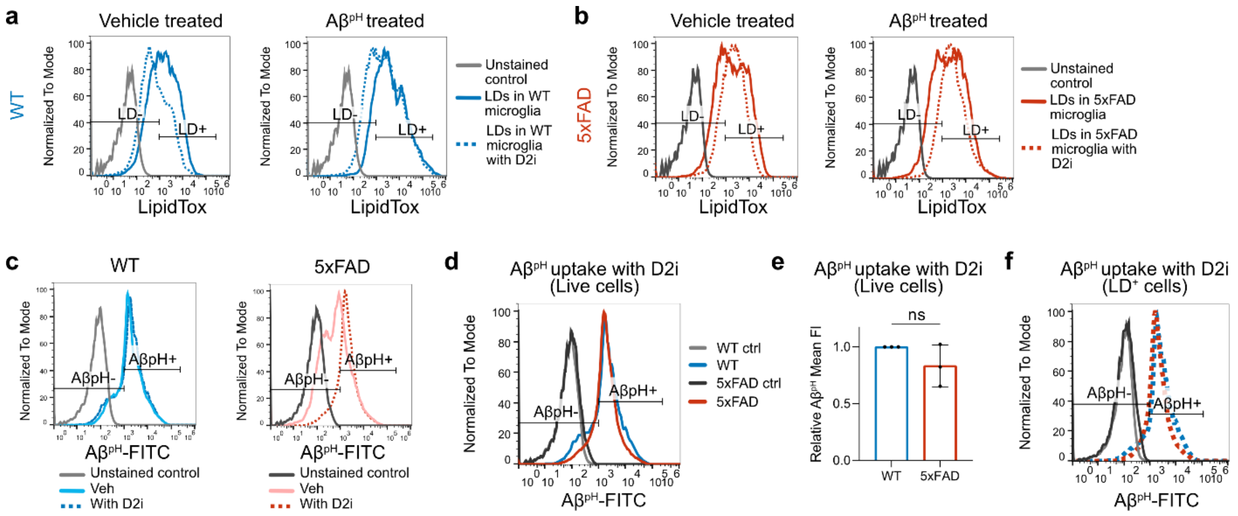


**Fig. S15. Effect of D2i treatment on LD formation and Aβ$^{pH}$ uptake in Aβ- and vehicle-treated 5xFAD and WT microglia.**

**a.** Overlay of representative histograms from vehicle-treated and Aβ$^{pH}$-treated microglia from 6-7-month-old female WT brains, with D2i treatment (dotted line) and without it (solid line). D2i treatment decreased fluorescence from LDs (indicative of LD load) in both vehicle- and Aβ$^{pH}$-treated microglia. **b.** Overlay of histograms from vehicle-treated and Aβ$^{pH}$-treated microglia from 6-7-month-old female 5xFAD brains, with D2i treatment (dotted line) and without it (solid line). D2i treatment did alter the LD content in vehicle- but not in Aβ$^{pH}$-treated microglia. **c.** Representative histograms comparing Aβ$^{pH}$ phagocytosis by live microglial cells from female WT (blue) or 5xFAD mice (red) with vehicle (solid lines) and D2i treatment (dotted lines) respectively. **d.** Representative histogram showing no difference in Aβ$^{pH}$ phagocytosis by live microglial cells with D2i treatment between female WT and 5xFAD mice. **e.** Quantification of Aβ$^{pH}$ uptake by live cells with D2i treatment. While a slight decrease in Aβ$^{pH}$ uptake was observed in D2i-treated 5xFAD microglia, it was not statistically significant compared to Aβ$^{pH}$ uptake by D2i-treated WT microglia. **f.** Histogram overlay comparing the signal of Aβ$^{pH}$ within LD$^{+}$ cells from D2i treated microglia from WT and 5xFAD brains. Quantification is shown in figure 4h. For **a-d, f**: Unstained controls for WT and 5xFAD shown as light grey and dark grey (solid) lines, respectively.

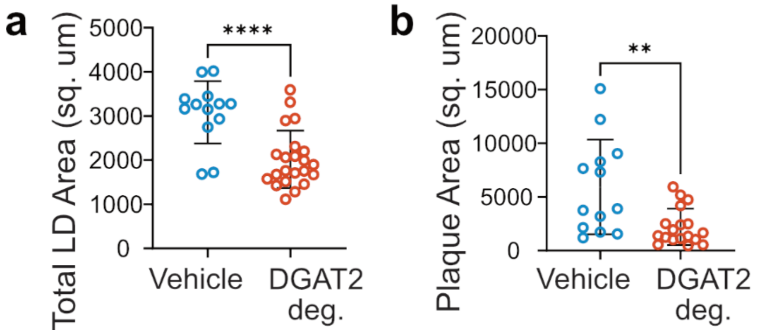


**Fig. S16. Effect of DGAT2 degrader on LD content and plaque deposition via delivery of a DGAT2 degrader in a subcutaneously-implanted osmotic pump in the lateral ventricles of 5xFAD brain.**

Quantification showing a significant decrease in **a.** total LD area and **b.** plaque area plotted for individual region of interest from all animals in the subiculum of 5xFAD brain treated with DGAT2 degrader vs. vehicle; ****$P$<0.0001, **$P$=0.0027. Data represent mean ± SD. Unpaired t-test, N=5 mice received DGAT2 degrader and N=3 mice received vehicle treatment.
